# Mapping the functional connectivity of ecosystem services supply across a regional landscape

**DOI:** 10.1101/2021.05.20.444939

**Authors:** Rachel D. Field, Lael Parrott

## Abstract

Sustainably managing multifunctional landscapes for production of multiple ecosystem services (ES) requires thorough understanding of the interactions between ES and the ecological processes that drive them. We build upon landscape connectivity theory to present a spatial approach for assessing functional connections between multiple ES at the landscape scale. We demonstrate application of the approach using existing ES supply mapping data for plant agriculture, waterflow regulation, and landscape aesthetics. The connections we observed between these three ES revealed high-value multifunctional linkages on the landscape that were not necessarily predictable from supply area mapping, nor from land use or land cover data. By providing spatial information on ES connectivity, our approach enables local and regional environmental planning and management that takes full consideration of the complex, multi-scale interactions between ecological processes, land use and land cover, and ecosystem service supply on a landscape.

## Introduction

The rapid, human-driven modification of wilderness is reshaping land cover distribution and disconnecting natural ecosystems, which has negatively impacted biodiversity and natural resources (Foley et al. 2005, Butchart et al. 2010, Foley et al. 2011). As these changes continue across the globe, ever-increasing demands are being placed on landscapes to deliver nature’s contributions to people, or ‘ecosystem services’ (ES; Carpenter et al. 2009). These juxtaposing forces highlight an urgent need for incorporating both biodiversity and ES in land use planning, with recent research calling specifically for consideration of landscape structure and connectivity in order to optimize environmental management objectives (Mitchell et al. 2013, Ekroos et al. 2014, Werling et al. 2014, Dobbs et al. 2014). The boom in ES research over the past several decades has improved our understanding of the ecological drivers underpinning the supply of ES, but more nuanced work is necessary to meaningfully manage ES provision and their interdependencies at the landscape scale (Kremen 2005, Tscharntke et al. 2005, Nicholson et al. 2009, Daily et al. 2009). Specifically, the supply of an ES is typically mapped within fixed areas (e.g., Tallis et al. 2008) without considering the potential relevance of ecological process flux across the landscape for supporting ES provisioning (e.g., Mitchell et al. 2013) and multi-ES relationships. By failing to represent the spatial and functional connectivity between supply areas in ES assessment, we ignore ecological processes that may be fundamental to the maintenance of ES supplies, run the risk of overlooking potentially critical areas in landscape-scale management, and miss opportunities for uniting divergent interest groups around the concept of multifunctional landscapes (i.e., those that provide multiple ES beyond those that are primarily managed; Power 2010). To optimize ES provisioning while minimizing potential negative effects on human well-being in the face of increased development pressures, it is critical to understand the dynamics of multi-ES supply (Lorilla et al. 2018).

Connectivity is a key attribute of landscape resilience and of ES in general (e.g., Bennett et al. 2021). A connected landscape facilitates the movement of energy, matter, organisms, seeds, pollinators, and people, thereby supporting several ecological processes that are critical for maintaining supplies of ES (Tscharntke et al. 2005, Kremen et al. 2007, Biggs et al. 2012, Mitchell et al. 2013, Pal et al. 2021). Conversely, a fragmented landscape exhibits decreased productivity, functional robustness, ecological richness (Leibold et al. 2004, Gonzalez et al. 2009, Simmonds et al. 2019, Melo et al. 2019), and increased vulnerability to further human modifications (Dutta et al. 2017, Xinxin et al. 2017, Liu et al. 2017, Chi et al. 2018). Smaller patches of habitat, biotic and/or abiotic supplies are not able to support as many species or as large populations relative to larger patches (Harper et al. 2005), and loss of landscape connectivity can hinder dispersal and migration of plants and animals (Fischer et al. 2007). Such deleterious effects on the wealth of biodiversity and natural capital can lead to declines in total ES supply and in the quantity and/or quality of flows to human beneficiaries (Mitchell et al. 2015, Pal et al. 2021). Pal et al. (2021) showed specifically that fragmentation, and not just diminishment of ES supply areas, contributed to significant declines in ES values over time. Landscape fragmentation impacts the supply of ES through altering the distribution and movement of the ecological elements, structures and processes underpinning the maintenance of natural capital (Mitchell et al. 2015). Mitchell et al. (2015) discuss how loss of connectivity can be a driver of interactions between multiple ES and can impact both the size and location of ES flows (Bagstad et al. 2013). Among key policy principles identified for enhancing ES resilience to disturbances and environmental changes is managing for connectivity among ES-related resources, species, and human actors, with specific focus on the strength and structure of these connections (Biggs et al. 2012). All this points to the importance of planning for connectivity in multifunctional landscapes (Phillips et al. 2015), while considering the potential for complex ecological process-based interactions among services, to successfully manage for the delivery of multiple ES (Dee et al. 2017).

In simple terms, planning for landscape connectivity typically focuses on habitat patches and movement corridors, whereas ES planning focuses on the areas of the landscape with the capacity to produce the services humans need to survive and thrive (Taylor et al. 1993, Egoh et al. 2008). Recent work calling explicitly for incorporating ES into connectivity research has taken the perspective of assessing how the characteristics of landscape connectivity (i.e., how a landscape promotes or hinders movement of matter and organisms), along with composition (i.e., quantities of land use and land cover, or ‘LULC’, types), and configuration (i.e., spatial pattern of LULC), might directly or indirectly impact ES provision and related ecological processes (Debinski and Holt 2000, Fahrig 2003, Gonzales et al. 2009, Mitchell et al. 2013). For example, adequate arrangement of adjacent natural habitat areas in agricultural landscapes can aid the movement of pollinators and pest predators to croplands, and thus promote delivery of these services (Priess et al. 2007, Ricketts et al. 2008, Fleischner 1994, Tscharntke and Brandl 2004; Kremen et al. 2007, Tallis and Polasky 2009, Power 2010, Lonsdorf et al. 2011). Habitat loss, fragmentation, and management strategies detrimental to related connectivity can have the opposite effect on pollinators (e.g., Kremen et al. 2002, Potts et al. 2010, Mitchell et al. 2013)., whereas increased connectivity in croplands can facilitate pest dispersal (Margosian et al. 2009). In terms of abiotic flows, connections between upstream and downstream freshwater sources can be important for maintaining quantity and quality of drinking water (Dodds and Oakes 2008, Bangash et al. 2013), and maintenance of the natural hydrologic regime stabilizes base flows and reduces flooding, thereby promoting waterflow regulation (Poff et al. 1997). Viewing the attributes of structural and functional connectivity in these ways, i.e., from the perspective of ES, helps to move beyond the idea of managing for spatially discrete ES supply areas and toward garnering better understanding of how we might manipulate certain connectivity elements (e.g., habitat patch area, isolation distance, movement corridors) to more effectively manage a landscape for individual ES (Mitchell et al. 2013). The above examples highlight that the ecological processes underpinning the supplies of certain ES directly influence the supplies of others, both when services co-occur in space and, sometimes, when they are produced in separate areas. Drivers behind multi-ES interactions, and the importance of such processes, are sometimes discussed in ES interaction research (e.g., Li et al. 2017, Alemu et al. 2020) but, to our knowledge, have not been explicitly delineated on the landscape in the context of multi-ES assessments. To truly understand the relevance of landscape connectivity to ES, we must ask: what and where are the mechanisms responsible for maintaining these connections and, consequently, supporting the production of multiple ES?

Since the seminal global appraisal of ecosystems and the ES they provide (MEA 2005), research that assesses the *interactions* between multiple services has increased exponentially (Agudelo et al. 2020; Appendix 1). However, research in this discipline commonly only considers services that co-occur in space (e.g., Queiroz et al. 2015), and assumes positively or negatively correlated ES to represent synergistic production or trade-offs, respectively (Tomscha and Gergel 2016, Agudelo et al. 2020). Such assessments are typically based on correlation coefficients of indicators aggregated within a geographic unit (e.g., watershed, municipality) or randomly sampled across a region (Anderson et al. 2009, Qin et al. 2015, Qiu and Turner 2015). These approaches to not directly evaluate interactions based on underlying ecological process theory nor do they allow for spatially discrete relationships to occur, i.e., they do not explicitly incorporate the mechanisms responsible for ES interactions, and they ignore how ES occurring in one area might have direct or indirect influence on ES in other areas. It has also been shown that simple spatial correlation analyses between pairs of ES are not necessarily a good predictor of how relationships between ES change over time (Mitchell et al. 2020), and that their interactions can vary across the LULC types found in heterogeneous landscapes (Li et al. 2017); thus, a better understanding of the processes that underpin the spatial patterns of ES is needed to improve the sustainable management of multifunctional landscapes (Mitchell et al. 2020). Recent research has visualized the spatial connectivity between ES supply areas by modelling the movement potential of species through high quality habitat corridors as a proxy for how biodiversity flow in general supports ES provisioning across the landscape (Peng et al. 2018). Still, this does not represent different functional connections between ES supply, and how the provisioning of one type of ES directly or indirectly effects the provisioning of another across a landscape. Further, a recent systematic review of studies that model interactions among multiple ES between 2005 and 2019 found that the vast majority of studies were conducted locally while relatively few studies were done at the regional scale (Agudelo et al. 2020). However, focus at the regional level may be most appropriate for reconciling the common scale mismatches between biophysical and socio-economic elements involved in sustainable ES management (e.g., Dalgaard et al. 2003, Cumming et al. 2006, Satake et al. 2008, Ingram et al. 2008), while minimizing practical issues with empirical mapping related to data gaps and indicator variability in areas larger than this (Verberg and Chen 2000; but see de Groot et al. (2010) for examples of variability in ecological scale relevance for specific ES). However, as this relates to *interactions* between different ES, small-scale observations may be masked at larger scales (Raudsepp-Hearne and Peterson 2016); therefore, incorporating local, grid-level data and analyses is important for providing meaningful information to planners (Haase et al. 2012).

In spite of the growing knowledge around the complex interactions and feedbacks between ES and the related suite of biotic and abiotic mechanisms and the cruciality of incorporating this into decision-making (Qiu and Turner 2013, Dee et al. 2017), spatial modelling of the diverse functional connections between multiple ES (e.g., Cui et al. 2012, Kolosz et al. 2018, Agudelo et al. 2020) from several broad ES categories at the regional scale remains limited (Field et al. 2017). Several approaches used in ecological connectivity studies to identify potential spatial linkages across a landscape are promising in their applicability multi-ES assessment. These include euclidean distances (Cressie et al. 1993), least-cost path analysis (LCP; Larkin et al. 2004), least-cost corridor (LCC; Singleton et al. 2002), circuit theory (McRae & Beier 2007), graph theory (Pinto & Keitt 2009), and network flow models (Phillips et al. 2008). All these approaches are potentially amenable to assessment of multi-ES interactions but, to date, we know of no studies that have applied such methods to map the process-driven interactions between the supplies of multiple ES in a regional context (Peng et al. 2018). Further, studies that have incorporated both landscape connectivity and ES concepts typically only focus on a single ES, are skewed toward specific types of provisioning (e.g., food) and regulating (e.g., pollination) services, and, to our knowledge, have not yet tested cultural services (Mitchell et al. 2013).

We present an approach to address the above research gaps, building on existing ES mapping and modelling and rooted in landscape connectivity theory, to demonstrate how the functional relationships between multiple ES can be represented in the context of connectivity planning across a regional heterogeneous landscape. We demonstrate our approach using existing grid-level data from a case study landscape in the southern interior of British Columbia, Canada, by mapping and assessing the connectivity between ES from three broad categories: provisioning (plant food agriculture), regulating (waterflow regulation), and cultural (landscape aesthetics; MEA 2005). Using these, we conceptualize ES supply areas as structural components, and the functional process links between these areas as configuration elements within a landscape connectivity framework. We base our approach on existing, and relatively straightforward, spatially co-occurring ES interaction and LCP corridor methods to present a first step toward representing functional connectivity between multiple ES. Our multi-step approach has three specific objectives: (1) to define the ecological process-based connectivity mechanisms between different types of ES supply; (2) to spatially map and quantify these connections while accounting for LULC heterogeneity; (3) to compare coverages of supply areas and functional connections across different types of LULC.

## Materials and methods

Our case study area spans the Okanagan region in British Columbia (BC), Canada, which we use to demonstrate a multi-ES connectivity mapping approach for informing landscape planning (Fig. 1). It is located in the south-central interior of BC, is Canada’s biodiversity hotspot and one of North America’s most endangered semi-arid ecoregions (Warman et al. 2004, Kerr and Cihlar 2004), has a highly diverse assemblage of land use types (see Caslys 2013), and covers 21,580 km^2^ from ∼276 to 2,774 masl. General LULC types in the region include, in decreasing order of area: forests (16,281 km^2^), grasslands (1,482 km^2^), natural parks (2,403 km^2^; NB: contains several of the other listed LULC categories), shrubs (1,349 km^2^), agricultural (842 km^2^), lakes (599 km^2^), urban residential (220 km^2^), rural residential (220 km^2^), wetlands (182 km^2^), rock/rubble (161 km^2^), exposed land (113 km^2^), manicured parks (45 km^2^), rivers (38 km^2^), commercial (23 km^2^), industrial (23 km^2^), urban institutional (16 km^2^), and reservoirs (6 km^2^; Field et al. 2017). The ES-related resources in the region are governed by over 80 institutions at federal, provincial, regional district, First Nations, and municipal levels. Our study area has strong engagement of regional government and non-government stakeholders, and an increasing demand for region-wide collaboration in land use planning to accommodate rapid rates of land development in response to a high rate of human population growth (Neale et al. 2007, Caslys 2013).

**Figure 1.**
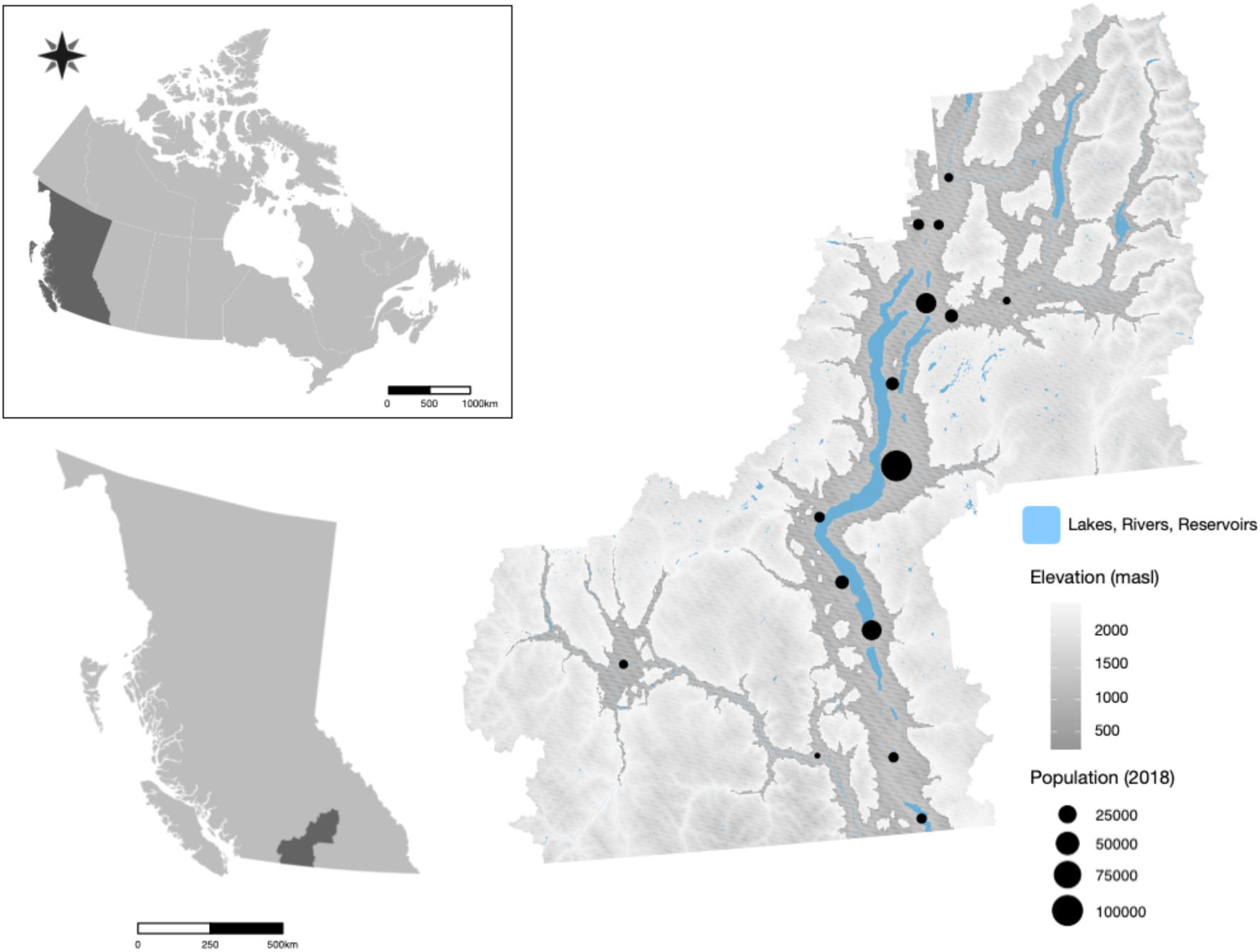
Location of the case study landscape in southern interior ‘Okanagan’ region of British Columbia, Canada. Major waterbodies, elevation (masl), and the most populous cities and towns in Okanagan regional districts are indicated.

We obtained spatial data for these ES from existing maps produced for our study area (Fig. 2; Field et al. 2017, Field 2021). Based on these data, we first established and valued *supply* area polygons - defined as spatially identifiable regions of higher-than-average-value supply potential - which serve as source and destination patches in a connectivity network (Appendix 1). We then developed a methodology to establish and value functional linkages, or connectivity, between supply areas. Functional connections were of two broad types: (1) *overlapping* links, which were areas where the supplies of two different types of ES occur in the same place, and there is an underlying process-based connection between them; and (2) *topographic* links, which were mapped based on the ecological processes that functionally connect the supplies of two ES areas separated in space. Lastly, we compared the coverage of top-value ES supplies and their linkages on the major LULC types found in the region.

**Figure 2.**
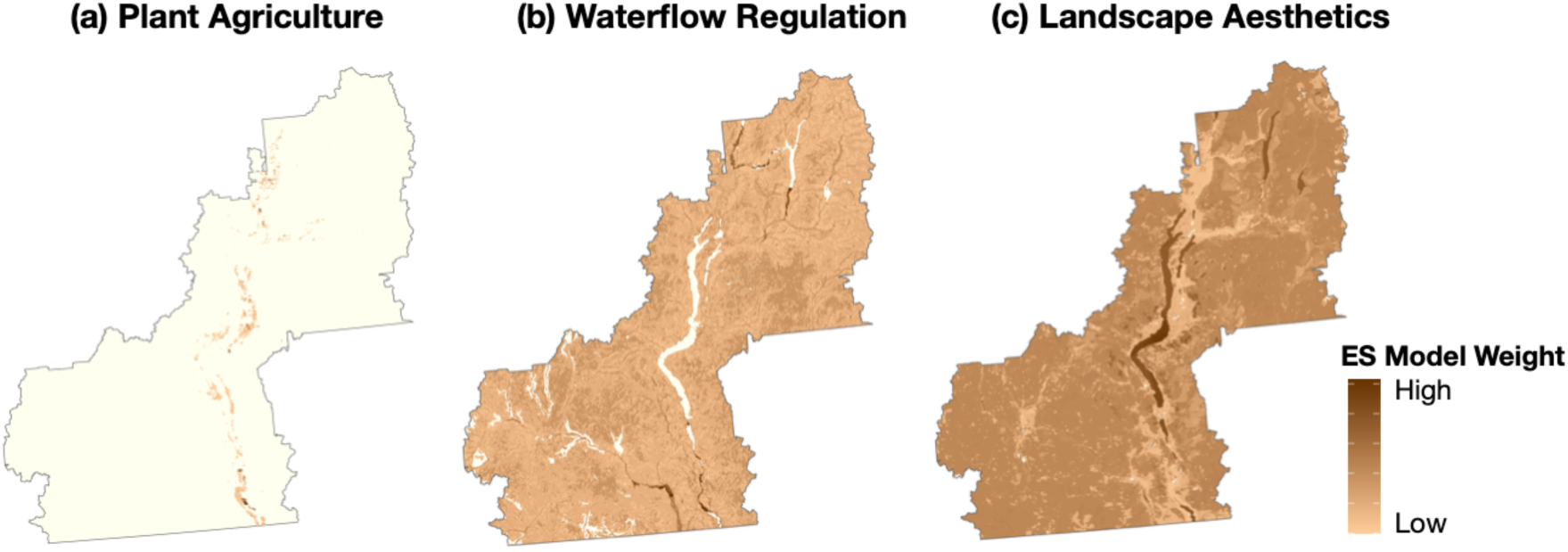
Maps showing the original, full-extent of distribution and weighting for ES supply areas in the case study landscape, including (a) plant food agriculture, (b) waterflow regulation, and (c) landscape aesthetics (Field et al. 2017).

### Mapping and valuing ES supply patches

To spatially partition the landscape into ES supply areas and establish the links between them, we developed a rationale based on interdisciplinary methods for assessing complex and connected natural systems (Bialonski et al. 2010). From the perspective of the landscapes’ capacity to provide ES, we defined subsystems as discrete areas with the greatest potential for providing ES supply, while the functional interdependencies between such areas were represented by spatial connections (also referred to herein as ‘links’ or ‘corridors’). We delineated ES supply areas based on approaches used in landscape connectivity and ES mapping studies: spatial polygons that represent high-value supply ES patch boundaries (e.g., Bangash et al. 2013); and the aggregation of immediately adjacent clusters of high-value supply spatial grid cells (e.g., Gardner 1999, Urban et al. 2009, Qiu and Turner 2013; Field and Parrott 2017). Any areas either lacking the potential for ES supply, or below a high-value supply threshold (details below), were represented as the landscape matrix through which ecological process-based connections between supply areas could flow (Field and Parrott 2017). In reality, such spatial interaction networks are dynamic through time (Boesing et al. 2020), though here we consider a static snapshot of the present state of ES supply in our study region in an effort to clearly illustrate real-world application of a novel approach for mapping the ecological relationships underpinning multiple types of ES supply.

For the purposes of demonstrating our concept, we used existing data on the spatial distribution of the ‘supply’ of three ES: (1) plant agriculture (‘PA’ herein; provisioning = products obtained from ecosystems); (2) waterflow regulation (‘WF’; regulating = abiotic and biotic processes that moderate natural phenomena); and landscape aesthetics (‘LA’; cultural = non-material characteristics that benefit human well-being; MEA 2005). Supply area maps were created using tools centred on ecological composition (biotic and abiotic elements), structure (e.g., topography, LULC distribution; Wallace 2007), processes (e.g., water infiltration, nutrient cycle, energy cycle; Lyons et al. 2005), and/or actual ES, and created primarily using existing and/or publicly available data. All data were resampled to ∼29 m x 29 m grid cells; therefore, ES models accounted for fine-scale heterogeneity of parameters across the landscape. We summarize the methods used for mapping the three (3) ES used in our study in Appendix 2 (Field 2021). Additional details are outlined in Field et al. (2017).

Studies that consider multiple ES have found that the distribution of at least one ES may be ubiquitous across a regional landscape (e.g., Queiroz et al. 2015), and/or isolated supply areas may be present within a non-ES-provisioning matrix (e.g., Qiu and Turner 2013; Fig. 2). Both situations were true for our landscape based on the ES selected, so for the purposes of creating supply areas, we chose to only retain areas with supply values above a top 50% threshold. This was because two of the ES types we selected, WF and LA, had near-ubiquitous spatial coverage with values ranging from very low to very high, and because management applications often are most interested in maintaining the highest-value provisioning areas (e.g., Turner et al. 2007). For PA, we subset the top 50%-valued polygons from the original mapping; for WF and LA, we subset the top 50%-valued raster cells of each of the regional ES maps, then converted these cells to single-part polygons based on aggregating adjacent cells within a diagonal raster cell width (∼29 m). Note that, due to the large file size of the WF data, the above was run separately for each sub-basin (n = 118) in our study area (see Appendix 3 for details). Aggregated areas became supply area polygons, and were valued based on the summed raster values therein, then normalized on a unit-less scale from 1 to 10000.

### Establishing functional connections between ES supplies

We define *ES connectivity* as areas on the landscape where one ES supply area influences the provisioning of another via underlying ecological processes. We identified spatial interactions between ES supply areas either as those that are connected through their overlap in space, or those that transverse the landscape through the relatively low value (i.e., sub-50% threshold) ES matrix. For these two cases respectively, we applied spatial overlay analysis (e.g., Qiu and Turner 2013), or identified flows using a stepwise procedure involving least-cost path (LCP) analyses akin to those applied in wildlife connectivity studies based on species movement and habitat attributes (Urban et al. 2009). Movement of organisms and matter across a landscape is often specifically defined in a single direction as a result of biophysical (e.g., waterflow, topography) or biological (e.g., movement from source to destination areas) realities, with multiple link types representing qualitatively unique flows that exist between patches (Zhang et al. 2007, Urban et al. 2009). For example, an area on the landscape producing multiple ES supply types may have functional links between ES of the same type in different locations, between different ES types in the same location, or with different ES types in different locations (Fig. 4).

**Figure 3.**
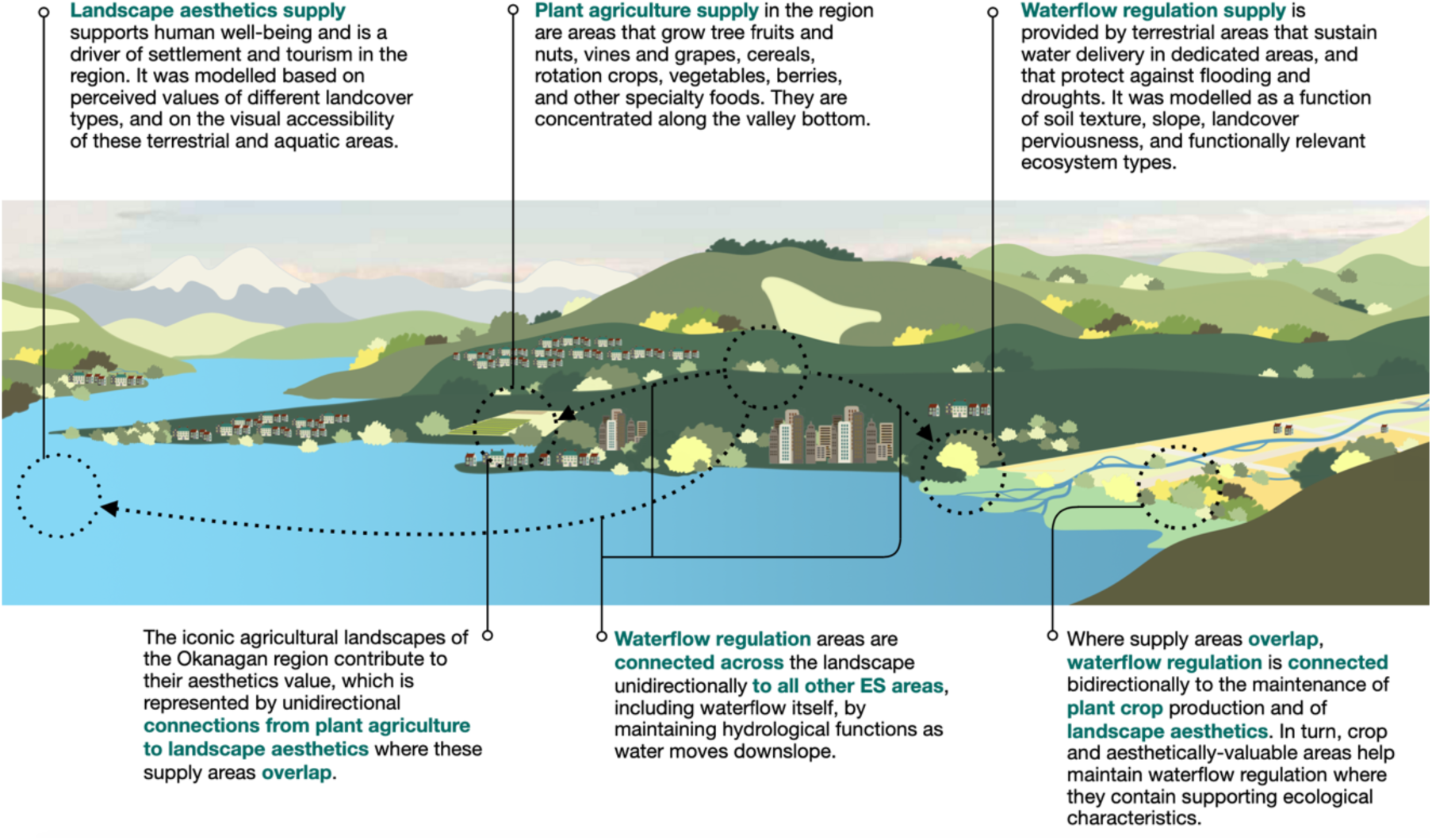
Schematic and definitions for ES supply areas and functional connections in the case study landscape.

**Figure 4.**
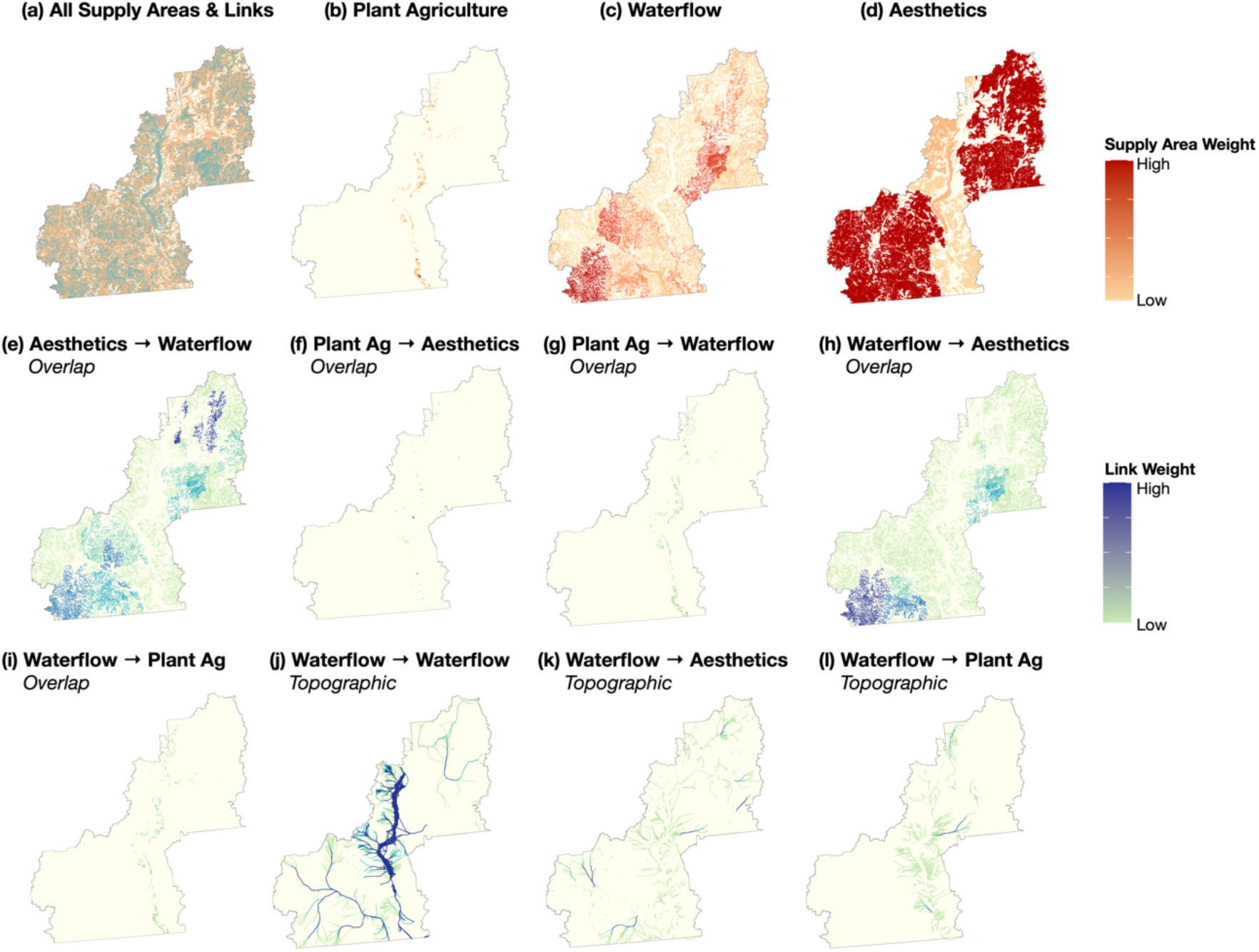
Conceptual schematic showing the functional connections between WF (purple), PA (green) and LA (orange) supply areas on a subset area of the case study landscape. All eight link types (topographic and overlapping) represented in our case study approach are differentiated by colour-coding, and weighted by thickness gradient. ES supply areas and links are nested within a non- or low-value landscape matrix (white area). Adapted from Ernstson et al. (2010) and Field and Parrott (2017).

For the three ES we considered in this study, we characterized eight (8) spatial link types by the directional, ecological process-based relationships between high-value ES supply areas. The rationale behind these connectivity mechanisms are summarized in Table 1. As connection model distribution and valuations were based on the original fine-scale supply area mapping, they also accounted for model parameter heterogeneity across the study area. We identified two high-level types of connections: overlapping (n = 5) and topographic (n = 3). *Overlapping* links were defined as areas where the supplies of two different types of ES occur in the same location on the landscape, and there is an underlying process-based connection between the two. For PA, the presence of vegetation crops can contribute to WF through providing a variety of beneficial ecological properties (e.g., soil texture, low-slope, high-perviousness, floodplains, riparian areas, and seasonally flooded fields; Power 2010), though the weight of this positive interaction may be higher if agricultural land was allowed to return to a natural vegetated state (Roa-García et al. 2011). PA also interacts with LA by providing farmland that is recognized as being aesthetically valuable (e.g., vineyards; Wagner and White 2009, Field et al. 2017) where these areas overlap. For WF, a direct positive influence stems from the spatial confluence of high-value WF areas on PA and LA supply areas through the maintenance of underlying hydrological processes where they co-occur (e.g., De Laney 1995, Nelson et al. 2009, Seavy et al. 2009). In the other direction, high-value terrestrial LA supply areas can be linked to WF areas through supportive ecological functions (e.g., pervious and water-retaining vegetated landscapes, floodplains in populated areas; Boyd and Banzhaff 2007, Van der Ploeg et al. 2010, Berkel and Verburg 2012, Carpenter et al. 2015; Table 1). We used the high-value ES supply area maps (Fig. 5b-c,d) to identify areas where each pair of ES overlapped (directionally) based on the above theory using a GIS-based clip procedure (see Appendix 4 for step-by-step details; Field 2021). The resulting single-part polygons of overlapping links represented the ecological processes connections between spatially co-occurring ES types (Fig. 5e-i).

**Figure 5.**
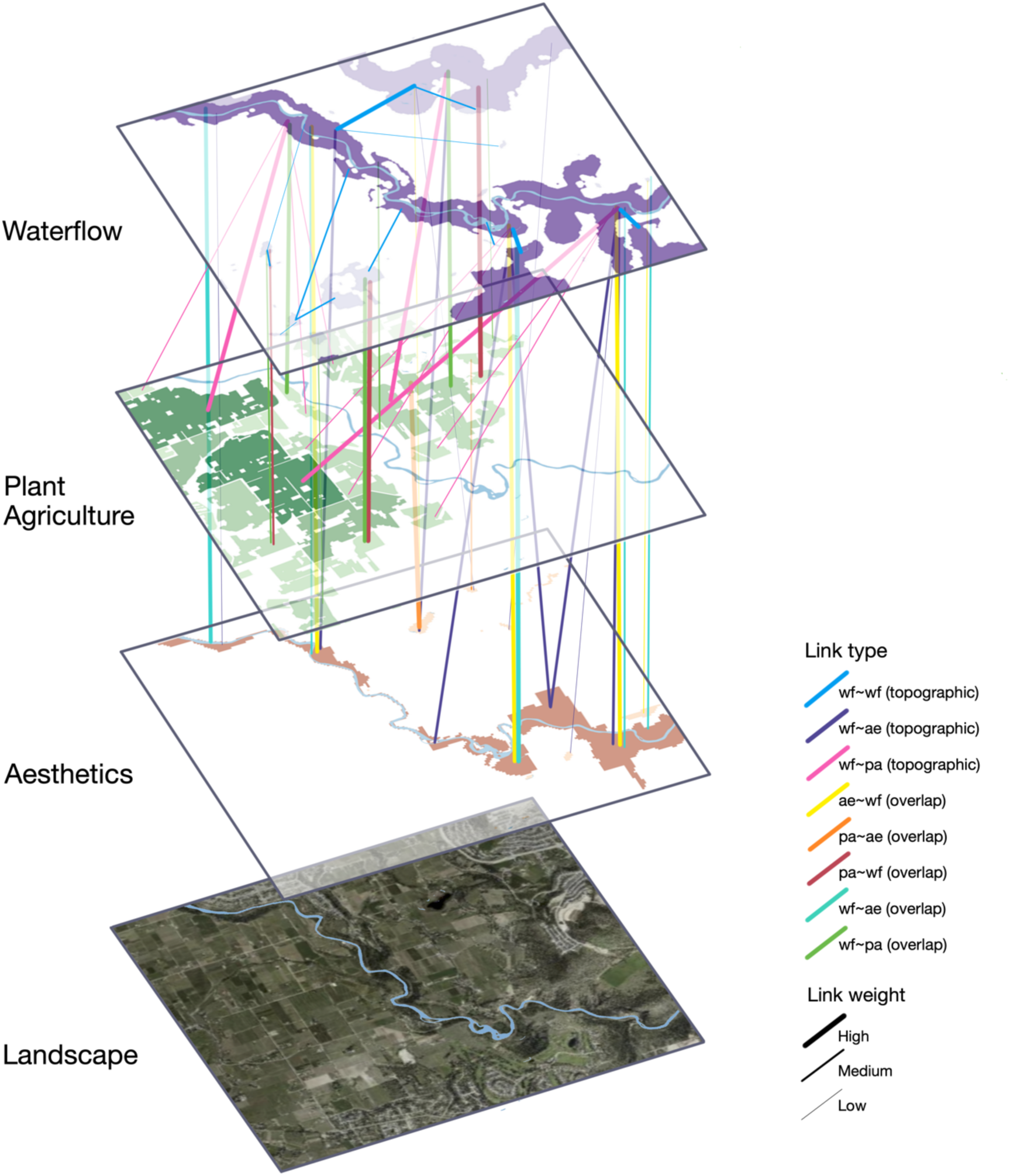
Distribution and weighting of top-50%-valued ES supply areas and functional connections on the case study landscape. Insets show (a) all top-value supply areas and links; top-value supply areas for (b) plant food agriculture (PA), (c) waterflow regulation (WF), and (d) landscape aesthetics (LA); overlapping connections from (e) LA to WF, (f) PA to LA, (g) PA to WF, (h) WF to LA, and (i) WF to PA; and topographic connections from (j) WF to WF, (k) WF to PA, and (l) WF to LA.

**Table 1.**
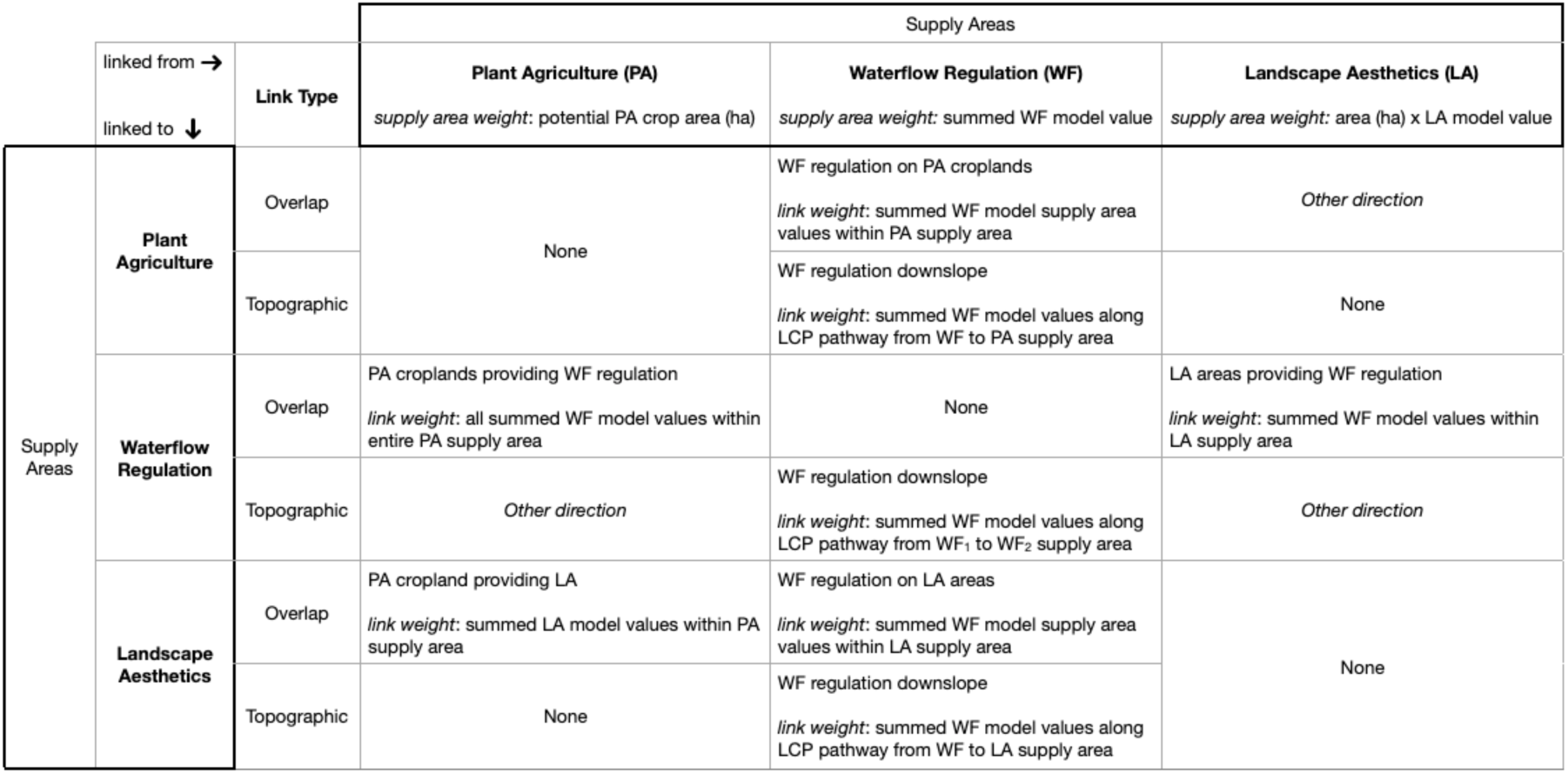
Rationale behind functional connection mechanisms, directionality, and weighting between top-value ES supply areas.

*Topographic* links were based on the ecological processes that functionally connect the supplies of two spatially separated ES areas across the landscape. Based on the three ES we considered, topographic connections always originate at a WF supply area, and represent the influence of upslope water regulation on the maintenance of the natural hydrologic processes that help support PA (e.g., crop growth and nutrient retention; De Laney 1995, Nelson et al. 2009), WF (e.g., natural baseline flow regulation; Nelson et al. 2009), and LA (e.g., maintenance of hydrology-dependent vegetation and aquatic features deemed to have high aesthetic value) supplies in downslope areas. We developed a stepwise procedure to create topographic links between ES supply areas. First, a separate least cost path (LCP) analysis was run for each WF supply area polygon to identify link corridors between these and other ES supply areas (Fig 6a). LCP analysis is a common method of mapping directional ecological corridors in landscape connectivity research, where the movement of an organism (or abiotic unit) is simulated across a resistance (cost) surface from a start to a destination point, and the lowest-accumulated resistance becomes the most likely path it will follow across a landscape (Beier et al. 2009). LCP can also be effectively used for hydrological flow models, where the algorithm seeks to minimize cumulative elevation along its path (e.g., Melles et al. 2011). The starting WF supply area polygon centroid was used as the ‘origin’ point for each associated LCP analysis. A single LCP ‘goal’ point was determined for each sub-basin by identifying the basin stream outlet (FLNRO 2017); the LCP goal point coordinates were identified as the intersect of this stream line feature and the valley-bottom line feature of the associated major watershed (Appendix 3; Field 2021). If multiple outlets were present in a sub-basin (e.g., Okanagan sub-basin ‘w11213’), the furthest downstream outlet line feature was used. LCP transition functions were built based on the assumption of downslope waterflow over a DEM surface, and allowed for connecting to a 16-cell neighbourhood to avoid paths being terminated based only on a single depression cell (van Etten 2017). Following this, we executed various procedures to produce topographic corridors between pairs of supply areas, which we summarize in Figure 6. This, along with approaches used to address other analytical nuances, are also discussed in Appendix 4.

**Figure 6.**
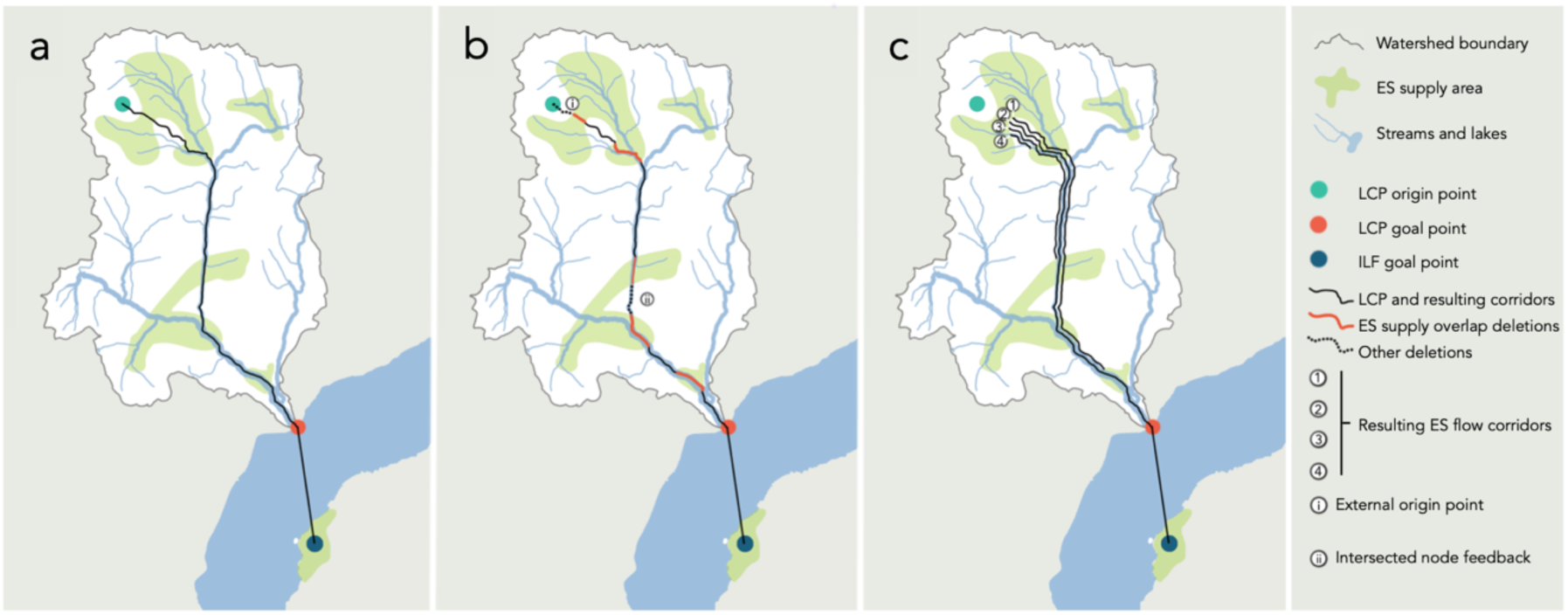
Schematic outlining steps for the creation of topographic ES corridors from each origin ES supply area to downslope supply areas. (a) An initial line feature resulting from a least cost path (LCP) analysis, i.e., from the origin to the goal point, amalgamated with a line from the goal point to a downstream influential landscape feature (ILF). (b) Types of LCP segment deletions addressed, including (red) segments overlapped by ES supply area polygons, (i) segments from origin points external to origin ES polygon, and (ii) segments flowing between two areas of an intersected (i.e., non-origin) ES supply area. (c) Resulting ES flow corridors after deletions, including feedbacks to origin ES supply area (4), flows to downslope ES supply areas (1) and (3), and flows to downstream ILF areas (2).

We then identified influential landscape features (ILFs) as additional WF polygons downstream of each sub-basin in the valley-bottom and associated with wetlands, floodplains, riparian areas, and/or seasonally flooded fields, which are functionally linked to upstream hydrological regulation. We connected ILFs to each upstream sub-basin outlet point, and individually merged these sub-basin lines with the LCPs for each WF supply area within that sub-basin (Fig. 6). Additionally, if a sub-basin flowed into a lake or reservoir, all ILF polygons immediately adjacent to that waterbody were included in the list of ‘downstream’ supply areas. Lastly, because the DEM raster resolution was approximately 20 m, some LCPs flowed outside sub-basin boundaries between the origin polygon (typically those close to a sub-basin boundary) and the goal point. Any nodes outside the sub-basin of interest that were overlapped by such LCPs were not considered to be true ‘intersections’ ’and such line segments were therefore excluded from within sub-basin links. However, such LCPs were still able to become connected to downstream ILFs.

### Assigning value to the functional connectivity between ES

In addition to spatially identifying connections between pairs of ES, we quantified the weight of these connections based on assumptions around the functional relationships between ES (e.g., Urban et al. 2009). We based valuations on the original ES provisioning maps, which assigned each raster cell in the map an ES value equivalent to results of the underlying models (Appendix 2; Field 2021; Field et al. 2017), and on the following assumptions. Since some of the ecological characteristics of both PA and AE areas can support the maintenance of WF supply, and high-value WF lands maintain the hydrological characteristics that help support PA provision and AE, we assumed that the value of these connections would approximate the available WF capacity. Therefore, the overlapping links between WF and PA, and between WF and LA, were weighted by the summed WF raster values therein. The plant food-providing agricultural areas of the Okanagan are prized by some for their beauty (Wagner and White 2009; though this subjective evaluation is complicated, see Wagner 2008), and we assumed the level of this significance to be equivalent to the underlying LA model value; therefore, such overlap links were unidirectionally weighted from PA to LA by the summed LA raster values therein. We assumed the contribution of upslope, high-value WF lands to the hydrologic maintenance of intersected downslope PA, WF and LA areas to be equivalent to the amount of flow regulation provided by landscape where water flows between these areas. Therefore, we quantified the weight of these unidirectional links by the cumulative value of all WF raster cells on the original map (i.e., not just top 50%; Fig X B original ES maps) traversed along corridors. All link weights were obtained by extracting summed raster cell values coincident with overlap link areas or with topographic link segments, then normalized on a unit-less scale from 1 to 10000. Additionally, raster overlay analysis was conducted, wherein cell values were summed across all eight link types to produce a weighted distribution map of all multi-ES connections for our entire study region. We acknowledge that alternative ecological process models could be used to produce more nuanced or accurate measures of link weightings (e.g., Cadotte et al. 2011). However, we chose to base our link quantification on high-level and readily calculable assumptions in an effort to provide simple, replicable, and easily-communicated metrics to inform applied decision-making for corridor, conservation, and protected area placement.

### Comparison with regional LULC

To compare the spatial coverage of supply areas and their linkages, and to aid in our assessment of potential uses of ES connectivity results for on-the-ground planning and management, we calculated the proportion of several high-level LULC categories intersected by each of the high-value supply areas and eight link types identified in the above analyses. We selected several LULC categories to provide both local and regional decision-makers additional information about where on the landscape ES connectivity is distributed, including forests, grasslands, shrubs, parks, aquatic areas, wetlands, rock and exposed land, agriculture, residential, and urban areas. We calculated the total area (ha and %) of LULC types covered by each link type, and the proportions of study area total LULC covered by each link.

ArcMap 10.7.1 (ESRI 2011) and R (version 3.6.2; R Development Core Team 2017) packages sp 1.4-5 (Pebesma and Bivand 2005, Bivand et al. 2013), sf 0.9-8 (Pebesma 2018), rgdal 1.5-23 (Bivand et al. 2015), raster 3.4-5 (Hijmans and van Etten 2012), rgeos 0.5-5 (Bivand et al. 2017), maptools 1.1-1 (Lewin-Koh et al. 2012), and stringr 1.4.0 (Wickham 2010) were used to build, assess and visualize the ES connectivity map. LCP analyses and subsequent stepwise link refinement were run using R package gdistance 1.3-6 (van Etten 2017). For transparency and reproducibility, data, R scripts and further details on our methodological procedures are available on the Open Science Framework (OSF; Field 2021; Appendix 5); limitations are discussed in Appendix 6.

## Results

### Distribution and values of ES supply areas

The top-valued 50% supply areas were distributed north-to-south across our study region (Fig. 5b-d). Plant foods are grown primarily in valley bottom areas in the Okanagan, and thus PA supply areas (n = 1,497) were concentrated in lower-elevation and population-dense regions with similar coverage to the original PA map (distribution of specific crop types detailed in Field et al. 2017). The highest-value PA supply areas were coincident with the largest farm parcels, present in the agriculture-rich areas of the south, north and east-central Okanagan. Given the extensive coverage of their original model results, top-value supply areas for both WF (n = 7,350) and LA (n = 5,262) were distributed fairly evenly across the entire study area. The highest-value WF supplies were associated with stream riparian areas in larger, partially-protected sub-basins of the southwest, and with riparian and wetland complexes in the central- and north-east. Our results suggest that the highest-value LA supplies were associated with large areas of upland forests, rivers, lakes, and protected parkland in the southwest and northeast, with relatively lower cumulative LA values in the more heavily-populated valley bottom. It is worth noting that, as our method of delineating distinct LA supply areas was based on the amalgamation of immediately adjacent raster cells, there were several large LA supply areas that may or may not be subjectively interpreted by human consumers as part of a single supply area. Issues with inherent subjectivity around LA mapping and assessment are common (e.g., van Zanten et al. 2016, see also Daniel et al. 2012), and this could lead to variable results in strength and physical location of cultural supply areas and their inter- and intra-ES linkages. Even nuances *within* a single cultural ES valuation method can lead to complex results; for example, tourist’s aesthetic appreciation of landscape features can differ from that of residents (Beza 2010). That said, the goal of this study is not to present the most accurate spatial representation of ES and their connections, but rather is to demonstrate a connectivity-based approach to evaluating multi-ES relationships. The original LA value distribution map is reproduced in Fig. 2c (Field et al. 2017).

### Distribution and values of functional connections between ES supplies

The spatial distribution and value of connections between overlapping ES were predictable based on the extents of supply area mapping and on the functional theory we applied to link weighting. Bi-directional overlap links between WF and LA (n = 9,363 in each direction) were distributed across the entire study area (Fig 5e,h). The highest-value links from LA to WF were associated with stream and lake riparian areas in both populated and remote valleys in the north, with riparian and wetland complexes in the central-east, and with remote stream and river riparian areas in the southwest. Similarly, the highest-value links from WF to LA were present in stream and river riparian areas in the southwest, and with stream riparian and wetland complexes in the central-east. For overlapping connections from PA to LA (n = 174), link distribution was sparse throughout the valley bottom and limited to croplands with high aesthetic value; primarily associated with vineyards and orchards (Fig. 5f). In terms of bi-directional overlap connections, the majority of PA supply areas were connected with WF regulation areas throughout the valley bottoms (WF to PA n = 1,220; PA to WF n = 1,320), with highly-weighted links typically associated with cultivated lands, fields, crop transitions, vineyards, and orchards near to (or containing) riparian, floodplain, and/or wetland areas (Fig. 5g,i).

Topographic links from high-value WF supplies to other ES supply areas revealed corridors variable in length and weight flowing across the landscape, sometimes linking ES supplies ∼200 kms apart (Fig. 3). Between pairs of spatially isolated WF areas, corridors (n = 484,602) approximated the location of watercourses (FLNRO 2017), as was expected due to the elevation-based LCP resistance surface used to simulate surface waterflow. The highest-value WF-WF corridors were observed through the large central Okanagan Lake system and several of its relatively low-order tributaries; in high-order valley-bottom rivers, streams and lakes in the southwest; and in the larger valley-bottom rivers of the northeast. These observations resulted from connections between WF supply areas and the ILFs that are prevalent next to valley-bottom aquatic areas. Flowing from WF to LA supply areas, corridors were scattered throughout the study area (n = 2,864), with the majority of links associated with the more populated valley-bottom areas in the central Okanagan basin (Appendix 3), and with the highest-value links in higher-order streams where sub-basins contained larger numbers of WF supply areas upstream of one or several LA supply areas. Connections to PA were only possible where farmlands were present within the sub-basin of the associated WF supply area, or downstream where farms were within ILF zones. Therefore, such corridors were concentrated in sub-basins along the central valley-bottom (n = 5,256), with particularly high weights in a northern agricultural valley used primarily for growing cereals and vegetables, in the largest sub-basin in the central Okanagan watershed primarily farmed for berries and tree fruits, and in a southern basin known for vineyards, tree fruits and vegetables. A general trend we observed for all topographic links was the co-occurrence of higher-value corridors with larger rivers and streams, rather than being associated with smaller headwater streams. This was a result of the culmination of overlapping corridors from several headwater WF areas in the lower-elevation stream valleys that had the largest number of supply areas for the related ES pair type.

When all link types were included on a map of accumulated weights, it highlighted expansive networks of high-value functional connectivity corridors between all three ES types and distributed across the entire landscape (Fig. 7). The highest-value link areas were found in low- and mid-elevation riparian areas across the landscape; in a mid-elevation wetland complex of the eastern-central region; in riparian and surface waterflow corridors associated with a large eastern-central sub-basin; and generally in areas where several (or all) of the eight link types co-occurred. Notably, the accumulation map revealed that several of the highest-value areas were not coincident with the highest-value on any of the individual link-type maps (Fig. 5), and were sometimes in relatively remote, higher-elevation areas.

**Figure 7.**
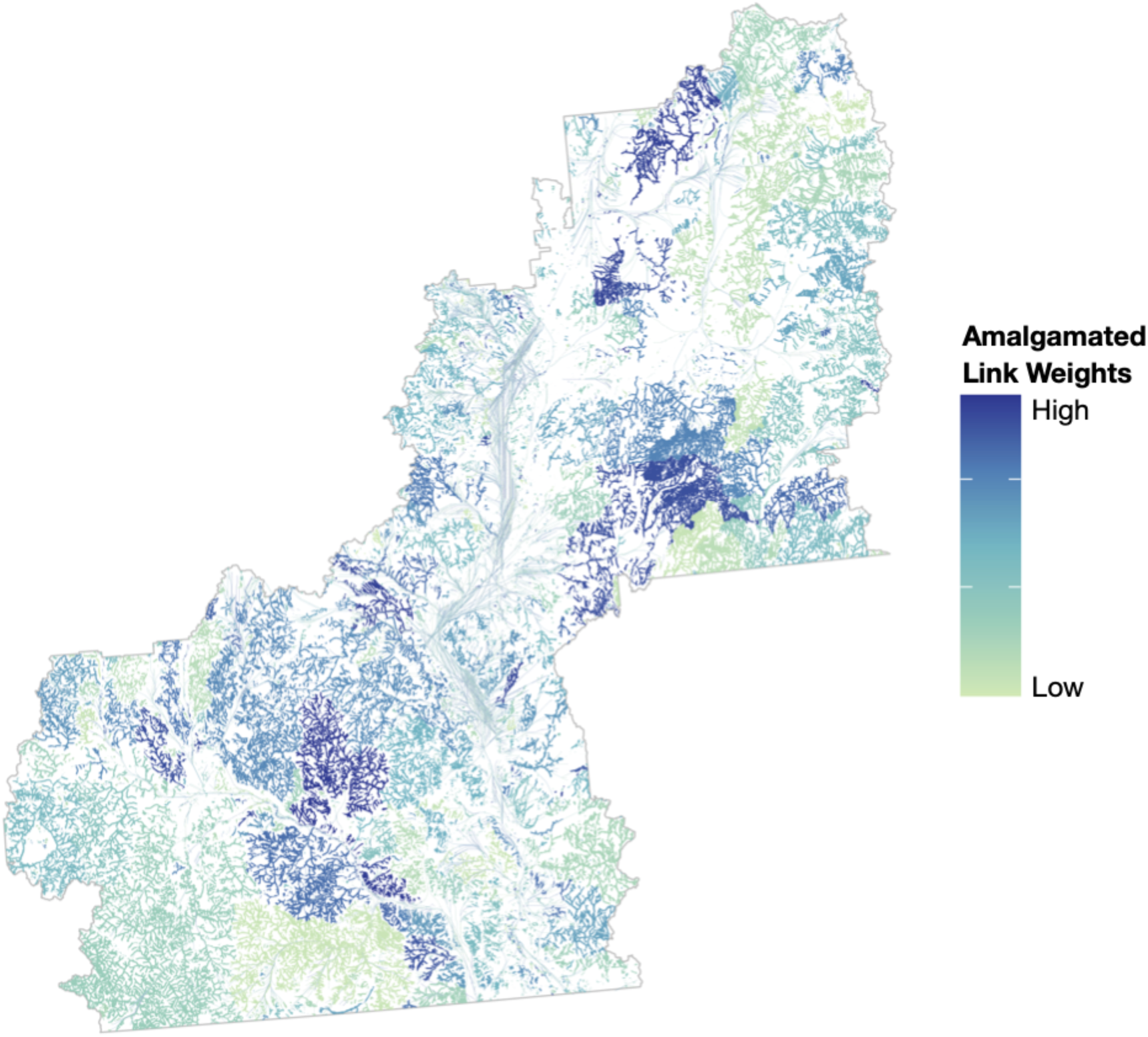
Distribution and weighting of link values amalgamated across all eight (8) overlapping and topographic link types across the case study landscape.

### Spatial coverage of supply area and linkages across LULC types

The distribution of ES supply areas and links were only in part determined by the underlying LULC types included in the original model parameters. In decreasing order of coverage of our study area (21,580 km^2^), the high-level LULC types are as follows: 76.7% forest; 11.9% park (NB: overlaps with forest, grassland, shrub, rock and exposed categories); 6.8% grassland; 5.8% shrub; 3.7% agriculture; 3.0% waterbodies (lakes, rivers, reservoirs); 1.6% residential; 1.3% rock and exposed land; 0.8% wetland; 0.3% urban; and 0.1% unknown (Appendix 7). For the subset top-50% ES areas, almost all PA supplies were, unsurprisingly, on agricultural lands (∼100.0%), but only covered 15.8% of all croplands in the region. Both WF and LA supplies were found mainly on forested lands (78.7% and 93.0%, respectively) and within parks (14.0% and 13.1%, respectively). Distribution of top-valued WF supplies covered large portions of most LULC types in the study area (19.7% to 50.6%; NB: 0% aquatic), including 99.6% of all mapped wetlands. Top-valued LA supplies spanned the majority of aquatic (98.8%), forested (81.9%), park (74.4%), and wetland (69.1%) LULC types (Fig. 8a-c).

**Figure 8.**
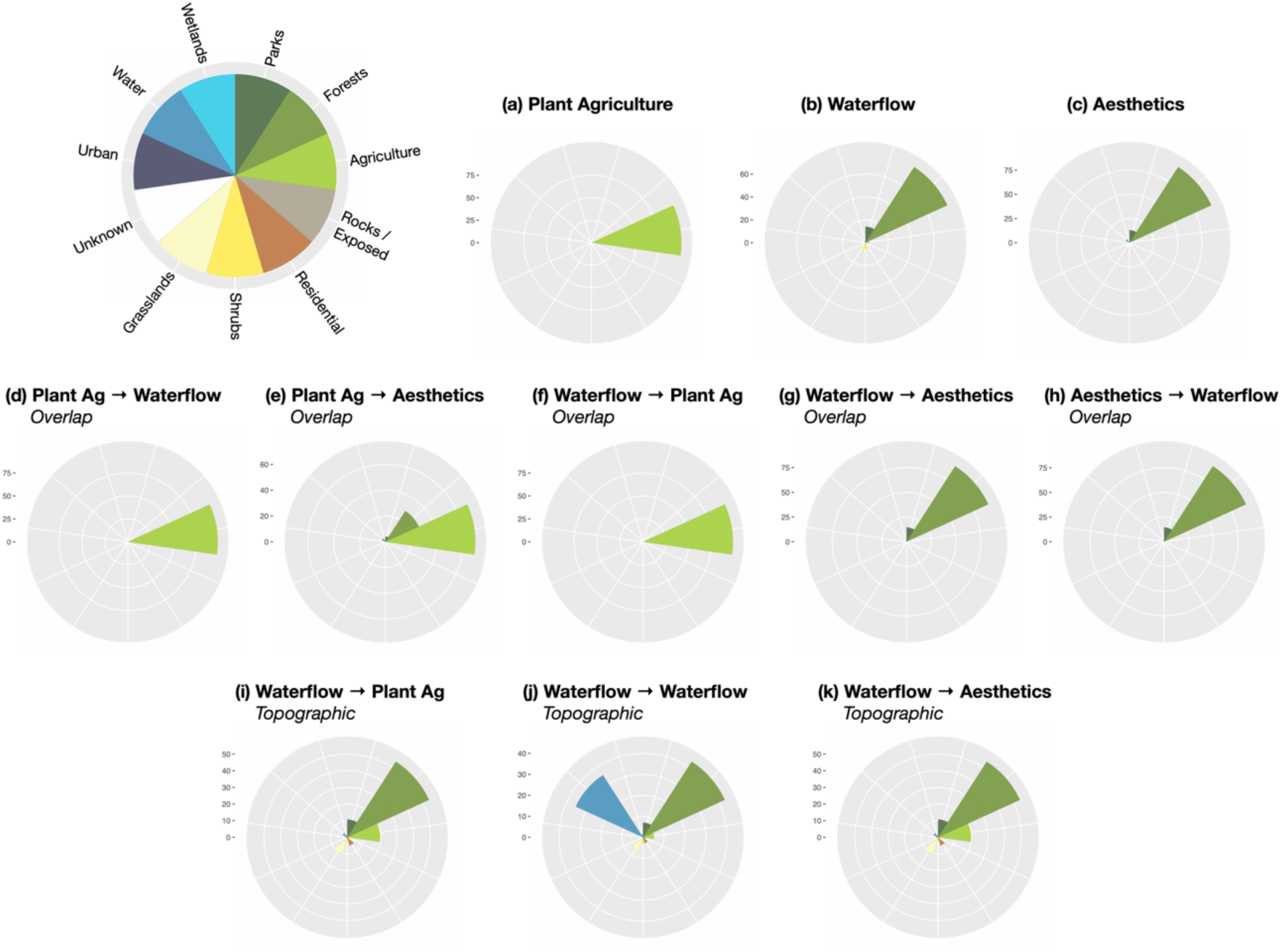
Proportion of major LULC types present within the extent of ES supply and link areas in the case study landscape. LULC types assessed are colour-coded and include forests, agriculture, rocks/exposed areas, residential areas, shrubs, grasslands, urban areas, water, wetlands, and areas with unknown use and/or cover. Insets show proportions in coxcombs for each top-value supply area, including (a) plant food agriculture (PA), (b) waterflow regulation (WF), and (c) landscape aesthetics (LA); overlapping connections from (d) LA to WF, (e) PA to LA, (f) PA to WF, (g) WF to LA, and (h) WF to PA; and topographic connections from (i) WF to WF, (j) WF to PA, and (k) WF to LA.

Similar trends in LULC coverage were observed for overlapping connections, with links from PA to LA found mainly on agricultural lands (69.7%) and in forests (28.5%). In both directions between PA and WF, connections were mainly on agricultural lands (both 98.0%), and covered 5.8% and 5.7% of all croplands in our study area from PA to WF and from WF to PA, respectively. In both directions between WF and LA, connections were mainly on forested land (90.8% and 91.1%, respectively) and in parks (14.7% and 14.6%, respectively). Moreover, these links covered large portions of all wetlands, parks and forests (from WF to LA 68.0%, 39.2 and 37.6%; from WF to LA 68.0%, 39.2 and 37.8%, respectively) in the study region (Fig. 8d-h).

For topographic corridors, we found that LULC coverage was less consistent with relevant ES supply area coverages. Corridors between different WF supply areas were found mainly in forested (42.9%) and aquatic (35.2%) areas, with more minor distribution in park (7.1%), grassland (6.9%), and agricultural (5.2%) LULC types. Notably, topographic WF corridors covered 24.3% of the entire aquatic areas found in our study region. From WF to PA, corridors mainly traversed forested areas (54.1%), followed by agricultural (19.8%), park (10.7%), grassland (10.6%), and residential (5.4%) LULC types. From WF to LA, corridors were mainly found in forested areas (57.3%), followed by parks (14.5%), agriculture (12.7%), and grasslands (10.2%; Fig. 8i-k). All LULC overlay analyses results are summarized in Appendix 8.

## Discussion

We applied a novel approach to mapping and modelling the functional connectivity between multiple types of ES across a regional landscape. By testing the application of this approach on three ES categories for a case study area, we identified and mapped eight link types connecting ES supply areas on the landscape. The results demonstrate the variety of ways categorically divergent ES can exhibit interdependencies related to their production potential, and the importance of considering these dependencies in land use planning for ecological connectivity.

### The case study: ES connectivity across a heterogeneous regional landscape

The connections we observed between ES revealed high-value multifunctional linkages on the landscape that were not necessarily predictable from supply area mapping. Across all link types we found heterogeneous distribution as well as spatially distinct areas of markedly higher value, or ‘hotspots’ of connectivity, relative to surrounding areas (e.g., Alemu et al. 2020). But one surprising observation is that the weighted amalgamation of all eight link types uncovered areas of high-value connectivity that were not present on any of the ES supply or pairwise link maps. This finding points to nuances that can be discovered when multiple ES *and* multiple linkage types are assessed together, and suggests that the spatial focus of planning for optimal service provisioning may shift when functional relationships between several ES are considered. Ultimately, such multifunctional areas represent possible conservation priorities that, if degraded or lost, may cause significant disruption of ES connectivity networks. Understanding the complexity of interactions between multiple ES has been highlighted as a critical challenge in planning for sustainable multifunctional landscapes in the face of changing environmental conditions and management interventions (Dee et al. 2017). A recent review of studies that have modelled interactions between multiple ES found that a large proportion did so from the perspective of co-occurring production synergies and trade-offs, but that the examination of flows, and the identification and quantification of explicit functional relationships remain largely unexplored (Agudelo et al. 2020). Ultimately, simultaneously modelling multiple ES continues to be difficult in part because of methodological inadequacies and the complexity of the ecological processes involved (Kolosz et al. 2018). Our approach provides a new framework that can help address these challenges.

From initiation points within WF supply areas, our modelling revealed several functional connections that operate over both short and long distances. Some of these topographic corridors extended over 200 km within the boundaries of our study area and, based on the underlying ecological process theory, also extend across the Canada-USA border to wetlands, riparian areas, seasonally-flooded agricultural fields, service supply areas along the extent of the Columbia River to the Pacific coast confluence between Washington and Oregon states, more than 1,100 km downstream from the originating supply areas in our study region. Similar long-range connectivity may be observed for other water-related ES (e.g., water provisioning, water quality), as both mean-annual water volume and water quality have been found to be heavily influenced by first-order headwater catchments, even in watersheds with large high-order rivers (Alexander et al. 2007, Freeman et al. 2007). Additionally, WF exhibits close- and long-range interactions with many other ES not modelled in our study. For example, water extraction and damming to take advantage of freshwater provisioning supplies alters natural hydrological regimes (e.g., Jackson et al. 2001); afforestation reduces peak and maintains base flows (e.g., Zhang et al. 2007, Power 2010), whereas deforestation destabilizes flows (Mäler et al. 2013); areas providing WF supply help to decrease pollution, flood-related turbidity, and residence time of chemicals in lakes (Burmil et al. 1999, Blackstock et al. 2001, Jackson et al. 2001, Bennett et al. 2009); certain pollination services can be facilitated by moving water (Biesmeijer et al. 2003); and some recreational activities are dependent on the maintenance of waterflow (e.g., fishing, kayaking; Burmil et al. 1999). Based on our observations of the potential for both short- and long-range functional connectivity, ES planning for other water-related services should also consider the potential impacts of management interventions on related services areas and management jurisdictions downstream.

Our study demonstrates that functional connections between ES often span several LULC categories, and that trends in dominant cover types may be unexpected relative to those associated with related supply areas. Certain areas or cover types are sometimes considered ‘hotspots ’for ES production, i.e., provide several different, often high-value, ES (e.g., Qiu and Turner 2013). For example, wetlands provide flood and flow control, storm protection, erosion control, groundwater supply, water quality maintenance, nutrient waste disposal, habitat to support fishing and hunting, natural materials, biodiversity, micro-climate stabilization, carbon sequestration, recreation, and aesthetic value (Brander et al. 2006). Agricultural lands can provide many ES beyond food for humans, such as habitat and food for pollinators, biological pest control (e.g., Loos et al. 2019), and tourism (e.g., Wagner and White 2009). We assessed the potential for LULC-associated connectivity hotspots in our region using LULC comparisons. Forested lands clearly stand out as being important for the regulating and cultural ES we investigated. Forests are often identified as hubs for maintaining regulating and cultural ES, including surface water quality, soil retention, carbon storage, and recreation (Matson et al. 1997, Brauman et al. 2007, Qiu and Turner 2013). Notably, although parks make up only 11.9% of the study area, they represent important landscapes for WF and LA supply and overlapping connectivity, and as flow corridors between all ES types we investigated. Both the above observations are likely driven by the suite of ecological processes present in complex forest, grassland and shrub ecosystems (e.g., vegetation-mediated infiltration, Mills and Fey 2004), and the contribution of wildlands and parks to aesthetics (Thompson 1995). From the perspective of functional connectivity, our study suggests a need to expand upon the ES ‘hotspot’ notion by considering that other LULC types beyond those associated with supply areas may be serving as critical corridors for interdependent ES. A clearly delineated example of this is the ecological process links between terrestrial and aquatic ecosystems. Areas of land adjacent to waterbodies are known to provide various regulation services in addition to WF, including erosion and water quality regulation through soil- and vegetation-mediated retention and filtration (Mills and Fey 2004). Whereas the model parameters we applied for WF preclude supply area coverage within any aquatic areas, the LULC proportions we observed within upland and downslope WF corridors transversed 24.4% of all surface waterbodies in the region and demonstrated that aquatic areas represent some of the most high-value linkages between different production areas for this ES. In addition, croplands proportionally represent the third-largest cover type in the corridors between upland WF and downslope LA supplies, with the majority of these corridors found in riparian zones, or on farms adjacent to wetlands and waterbodies. The synergistic association of WF and PA supply areas has been observed in other ES interaction studies (e.g., Qiu and Turner 2013), and stems from crops’, especially deep-rooted perennials, ability to provide a variety of hydrological benefits including increased water infiltration and recharge, reduced runoff, and mitigation of peak flows (Dabney 1998, Tilman et al. 2002, Brauman et al. 2007, Power 2010). These observations have implications for ecosystem- and habitat-based management programs as LULC types are often imposed as boundaries for interventions and/or institutions (e.g., BC Ministry of Agriculture). Especially in heterogeneous regional landscapes, our results point to potential for increased need for cross-jurisdictional collaboration when planning for functional connectivity in the optimization of multiple ES.

Ecological degradation and climate change-induced aridity is increasing across the globe. This has detrimental impacts on the structure and function of dryland ecosystems, which are characteristics strongly related to the ability of these systems to produce essential ES (Middleton and Sternberg 2013, Huang et al. 2015). The high interconnectivity of natural biotic and abiotic attributes can lead to cascading changes. Single ecological attributes can have negative impacts on several others (Schlesinger et al. 1990, Reynolds et al. 2007, Maestre et al. 2016); can exhibit sudden non-linear responses to increased aridity; and can cross sequential, multi-phase thresholds of ecosystem change. This can further be complicated by interactions with topographic-and land use-related factors. Climate forecasts predict that up to 28.6% of current drylands will cross one of the three sequential aridity-induced degradation thresholds by 2100, broadly characterized by vegetation decline, soil disruption, and systemic breakdown (Berdugo et al. 2020). This is relevant to our study region as it comprises the northern extent of the semi-arid North American Great Basin ecoregion, with high-elevation mountains flanking its eastern and western boundaries. This topography leads to the Okanagan region’s function as a critical movement corridor for north-south species migrations (Transboundary Connectivity Group 2016), and it likely exhibits similar pinch-point connectivity dynamics for topography-influenced ES, as we observed for the WF corridors modelled our study. The potential for imminent and irreversible degradation of ecological processes in dryland systems calls attention to the desperate need for understanding potential implications of these effects on multiple ES and their interdependencies. Our study is a jumping-off point for a more thorough evaluation of future aridity risks to ES relationships in the Okanagan and other dryland regions.

### The conceptual shift: from correlative interactions to functional connectivity

Designing landscapes for optimizing delivery of ES, while minimizing ecosystem degradation, requires thorough understanding of the interactions between multiple services and the ecological processes that drive them (Fu et al. 2013, Agudelo et al. 2020). Key guiding paradigms for this, and indeed landscape ecology in general, are to understand the relationships between spatial patterns and ecological processes, and to consider not only individual components (e.g., a single ES or supply area) but their combined effects (Wallace 2007, Nassauer and Opdam 2008, Fu et al. 2013). Viewing ES in terms of their spatial interconnectivity is especially relevant to regulating and supporting services that are known to be connected to the supply of many others; e.g., water flow is essential to the production of water for drinking, irrigation, and industry; plant and animal food production; regulating microclimates; and, as we exemplify in this study, landscape aesthetics (Fu et al. 2013).

Areas of ES supply are not necessarily spatially congruent with the discrete structural components traditionally considered in landscape connectivity frameworks (e.g., habitat patches, specific LULC types); therefore, linkages between ES are also unlikely to be coincident with these components (e.g., movement of organisms and matter; Brooks 2003). For example, the global benefit of carbon storage and sequestration depends only on the quantity of natural landcover, and not the spatial arrangement of patches (Mitchell et al. 2015). Although protected areas and intact habitat patches are important spaces for some of the plants, animals, and abiotic ecosystem components responsible for providing ES, provisioning and flows are not bound by human-defined reserve areas, and many ES are produced completely by and interact with one-another in human-modified landscapes (Schröter et al. 2019). Further, connectivity of certain ES will be limited by distance thresholds and/or human or ecological barriers to the flow of ecological processes. For example, crop production can benefit from interspersion of natural habitat throughout agricultural landscapes, which can increase pollination and pest control services delivery from species that can only move limited distances from their habitat patches (Tscharntke et al. 2005). There may be spatial congruency between existing wildlife movement corridors and certain regulating services, especially those that depend on the movement of organisms for their delivery, e.g., pollinators, disease control, pests and their predators, seed dispersal; Kremen et al. 2007), which suggests that there may be opportunities for win-win conservation initiatives for wildlife and ES together. Our approach can be used to explore this possibility, and to explicitly map and assesses the mechanisms behind distance-threshold-mediated and cross-landscape ES interactions in general. Outcomes of such work can be readily incorporated into connectivity-based management for these and other complex interactions that operate across heterogeneous landscapes.

Our approach reveals cross-landscape connectivity processes that represent important drivers of ES production, and are undetectable with traditional approaches for identifying ES synergies and trade-offs (e.g., Qui and Turner 2013, Su and Fu 2013, Tomscha and Gergel 2016). It can be used to represent several different types of functional connections, e.g., between different ES that occupy the same space, and abiotic movement from one ES supply area to another across the landscape. Identification of links between spatially co-occurring supply areas is similar to a representation of paired ES ‘interactions’, a concept for identifying synergies and trade-offs among services, as well as identifying groups of services that repeatedly occur together across a landscape (i.e., ‘ES bundles’; Bennett et al. 2009). Co-occurring ES typically arise as a result of common drivers or direct interactions among ES. For example, fertilization can drive-up crop yield while simultaneously decreasing water quality. Such services can interact through the same underlying ecological function (e.g., forests can have high recreational and aesthetic value and also provide water flow regulation). Further, ES in the same ‘neighbourhood ’can interact with one another (e.g., pollinator habitat adjacent to crop lands can increase crop productivity; Cord et al. 2019). Our methods take a closer look at the concept of ES interactions by explicitly representing the mechanisms behind the co-occurrence ES in the delineation and valuation of these areas (Bennett et al. 2009). Investigation of interaction mechanisms with respect to multi-ES assessment has been highlighted as a crucial step toward providing more rigorous information to inform the management of multifunctional landscapes (Alemu et al. 2020), and our study is one of the few to provide this information at the regional scale (Agudelo et al. 2020).

### Practical implications

Landscape planning typically involves considering the spatial layout of different, often incompatible, land use types that ultimately influence connectivity (Phalan et al. 2011). From an ecological connectivity perspective, such guidelines may include the maintenance of biodiversity to support ecosystem function, and thereby ES production and resilience, by including an assemblage of native habitat patches, corridors, stepping stones, and sensitive habitat buffers within a structurally complex matrix, with more focused management strategies for species and/or functional groups (Fischer et al. 2006). Notwithstanding that biodiversity can have complex, non-linear relationships with ES production (Fu et al. 2013) and is not necessarily directly related to multifunctionality (Birkhoffer et al. 2018), one drawback of this approach is that a disconnect emerges between the general maintenance of ecological connectivity and the benefits realized by humans. Recent conceptual frameworks have called for research that uncovers a more nuanced understanding of how landscape structure impacts the provision of *multiple* ES, how abiotic flows impact ES provision, and which services are more influenced by landscape connectivity (e.g., Mitchell et al. 2013, Mitchell et al. 2015). However, by not explicitly considering how ecological processes result in the production of ES and how these ecological processes are connected (Fu et al. 2013), there is risk of spatial incongruence between traditional connectivity planning priority areas and actual ES supply and flows across the landscape. Our approach helps to address this risk by enabling spatially explicit identification of high-value multi-ES links across the landscape, which can be overlaid and compared with other connectivity priorities (e.g., wildlife corridors).

### Opportunities for future work

Our case study maps and quantifies relationships between ES at a snapshot in time. However, modifications of natural landcover can change the number, size, shape, isolation, and distribution of ecological patches across the landscape and their proximity to human beneficiaries, all of which may lead to positive, negative or neutral impacts on ES supply and flow (Mitchell et al. 2015). For example, agricultural intensification tends to negatively impact large pollinators more than small ones, the former of which are more efficient crop pollinators (Suding et al. 2008). Both types of pollinators could be represented in a multi-ES network, and potential impacts of land use change could be addressed using modelling of future scenarios (e.g., Redhead et al. 2020). In the face of climate change, increases in dryland aridity causes grasslands and savannahs to metamorphose into shrublands as the latter grow better in sandy, nutrient-poor soils (D’Odorico and Okin 2012, Phillips et al. 2019). In our study area specifically, such a shift would have implications for ES coverage and value through the dependency of model variables on underlying vegetation characteristics (Field et al. 2017), and thus influence future ES production and connectivity. The ultimate impacts of landscape changes on ES are dependent on the structure and quantity of such changes, and on the biophysical process, ecosystem functions, species, and human activities driving the ES supply of interest, as well as the flows to and demands of human beneficiaries (Mitchell et al. 2015). Even if human development does not greatly diminish the quantity of natural landcover, it can still have far-reaching impacts on ES supply. For example, the prolific damming of the majority of major river systems has reduced the flow of water and the use of these systems as corridors for human movement, and changed natural patterns of water provisioning, water quality regulation, and recreation opportunities (Loomis 2002, Whittaker and Shelby 2002, Nilsson et al. 2005). Further, it has been shown that spatial correlations between pairs of ES can exhibit inter-annual variability (e.g., Renard et al. 2015, Li et al. 2017), and that snapshots in time are not good predictors of how their relationships may change over time (e.g., Mitchell et al. 2020). Future studies could use the ES connectivity framework presented here to assess how changes in LULC ultimately have cascading impacts on multiple ES across a landscape (Bagstad et al. 2013, Grêt-Regamey et al. 2017, Rieb et al. 2017), which can practically be achieved by incorporating seasonal and inter-annual variations in ES supply, demand and functional connectivity (e.g., increases in fresh water provisioning during dry months; Field and Parrott 2017). The application of our framework along with scenario modelling and/or historical data could help us to move beyond examining how natural and anthropogenic drivers of change impact local provisioning of an individual ES or trade-offs between multiple ES (e.g., Li et al. 2017, Schroter et al. 2019), to how such isolated drivers might have far-reaching, cumulative impacts at multiple spatial and temporal scales.

We acknowledge that several other methods exist for identifying and evaluating corridors across a landscape, including least cost corridors (Singleton et al. 2002), circuit theory (McRae and Beier 2007, McRae et al. 2008), graph theory (Fall et al. 2007, Pinto and Keith 2009, Rayfield et al. 2011), networks (Phillips et al. 2008; Parks et al. 2013), and deterministic eight models (Mark 1984). We chose to employ only LCP analysis mainly because the topographic ES flows in our study all originated at WF supply areas, corridors all were to represent the ecological process of water flowing downslope, and because LCP has been shown to be a valid method for approximating drainage networks while being capable of overcoming issues around topographic depressions (Melles et al. 2011). Therefore, a DEM-driven model representing water moving downslope was deemed the most appropriate for these types of ES connections in our study region. Further, our aim was to provide relatively simple representations of corridors between supply areas to support the primary goal of this paper, i.e., to demonstrate a novel approach for conceptualizing how the provisioning of ES are functionally connected across a landscape. It is important to highlight that our method of valuing topographic links corresponds to the accumulated weights of relevant ecological processes along the entire length of a corridor. We do not attempt to quantify the potential strength of contribution the upstream supply area exerts on an individual downstream supply area (i.e., contribution to maintaining the downstream area’s potential for producing baseline service supply). A promising area of future research for addressing the latter distinction will be through the use of network theoretic methods (discussed below), which can employ patch (e.g., supply area or ‘node’) and link valuations, as well as viewing these in the context of the entire network, to calculate metrics of the overall influence of an individual patch (e.g., Field and Parrott 2017). In summary, future research could compare and validate alternative spatial corridor mapping and valuing approaches (e.g., Melles et al. 2011) for predicting process-based movement between WF and other ES types.

Landscape connectivity studies have assumed that larger ecological patches are more robust and valuable to a network (e.g., Pal et al. 2021). However, especially for multiple ES that span a variety of habitat types, for those influenced by below-ground processes (e.g., soil texture - WF, Brauman et al. 2007; sub-surface carbon stock - climate regulation, Friess 2016), and for those dependent on landscape features that are typically independent of ‘patch’ arrangement (e.g., topography - WF, Crossman et al. 2013), additional methods should be explored to identify alternative methods of delineating and valuing patches for ES connectivity. Spatial network (graph) theory has been used to assess the movement of individual ES across the landscape (e.g., Janssen et al. 2006, Fortuna et al. 2006, Heckmann and Schwanghart 2013, Phillips 2013, Peron et al. 2014) but, although it has been called for (e.g., Bohan et al. 2013, Hines 2015, Quintessence Consortium et al. 2016, Dee et al. 2017), studies on real-world networks with different types of linkages between multiple types of ES, especially relating to all the broad categories of provisioning, regulating and cultural services, are still lacking (Field and Parrott 2017). In spatial network analysis, landscape components (e.g., habitat patches) are represented as ‘nodes’, and connections (e.g., habitat corridors, species dispersal) form the ‘links’ between them. Once built, network metrics can be used to evaluate a variety of characteristics including the strength of connectivity, most likely flow routes, relative contribution of individual nodes to overall connectivity, flow efficiency, node and link vulnerability to disturbances, etc. (e.g., Urban et al., 2009; Heckmann et al., 2015). Network analyses have been applied to well-described species-specific mutualistic interactions as they relate to indicators of ES supply (e.g., pollinator and seed-disperser relationships with plants, Gilarranz et al. 2011). A recent single ES and species-specific case study on the relationship between river red gum tree and periodic flooding (i.e., WF) used a conceptual network framework based on the functional drivers and feedbacks between abiotic, biotic, and social system components to guide management of the system (Dee et al. 2017). However, we know of no study that has considered combined biotic and abiotic contributors, both of which are necessary to build an accurate picture of ES supply across real world landscapes. Our approach provides a framework for network-based ES assessment by spatially representing interactions and feedbacks between multiple services from three broad categories. It is also amenable to situations where there are several distinct link mechanisms (i.e., multiplex network; Horvát and Zweig 2012), resulting in positive and negative interactions between the same ES types, all of which can be analyzed in a spatial network framework. For example, links between plant agriculture and water quality regulation areas could be driven both by positive unidirectional flows from (especially perennial) croplands providing water filtration services, and by negative unidirectional flows from croplands that use fertilizers and/or pesticides. The next step will be to use spatial network analyses to assess our multi-ES framework from the perspective of supply area ‘nodes’ and ES connection ‘links’, which will allow for the incorporation of both qualitative and quantitative social-ecological data and the evaluation of metrics related to functionality of supply areas, their connections, and the entire ES network (e.g., Field and Parrott 2017).

## Conclusions

Our study provides a new approach for the assessment of multiple ES and provides important information on the spatial interconnectivity of a variety of divergent types of ES across a diverse temperate landscape in southern interior British Columbia. We are confident that providing a tool for visualization of multiple ES will help address several ongoing challenges: increase awareness and understanding of how dependent humans are on nature; highlight a need to maintain landscape connectivity to support ecological functioning; advance the interdisciplinary science around the ES concept; and help move toward incorporating this science into management of natural capital (Guerry et al. 2015). As the ES concept continues to be developed and refined, considering how ES operate within the context of interconnected, complex social-ecological systems will help improve our ability to meaningfully incorporate multiple ES into decision-making and planning at the landscape scale. Overall, our methods not only allow for the explicit incorporation of the current knowledge of the ecological processes driving linkages between multiple ES, but they also provide decision makers mapping tools that show where these connections occur on the landscape and how valuable they are to ES production potential. Thus, our approach can help guide planners in predicting how intervention(s) in specific location(s) are likely to have synergistic or antagonistic impacts on ES supply areas in other, sometimes distant places.

## Acknowledgements

We received constructive feedback from L. Hooker, J. Janmaat and J. Pither. Funding for this manuscript was provided by the Regional District of Central Okanagan, the Okanagan Collaborative Conservation Program, a Discovery Grant from the Natural Sciences and Engineering Research Council of Canada (NSERC) awarded to L.P., and by a Canada Graduate Scholarship from NSERC awarded to R.D.F.

## Appendices

## Appendix 1 Background information

### ES interactions

Other work has defined ‘interactions’ as value-based synergies (increase in supply quality and/or quantity of one ES results in supply increase of another), trade-offs (increase in supply of one ES results in decrease of another; Bennett et al. 2009), bundles (groups of ES that co-occur repeatedly across a landscape, typically linked to co-variation in LULC types; Raudsepp-Hearne et al. 2010, Lee and Lautenbach 2016), or flows (ES interactions from the perspective of beneficiaries) between ES that occur over the same space and time (Agudelo et al. 2020).

### ES supply

We note that, although we use the term ‘supply’ to refer to the portion of the ES provisioning delivery chain on which we are focused, two of the ES we have selected can be conceptualized as spanning both the supply and ‘flow’ aspects of this chain. ES supply refers to the ecological good(s) and service(s) produced by a natural or man-made area on the landscape (Potschin et al. 2016), whereas ES flow represents human access to ES supplies, i.e., the transfer of a good and/or service from a supply to a benefit area or actor for use (Villamagna et al. 2013, Schröter et al. 2018, Schirpke et al. 2019, Vallecillo et al. 2019). For agriculture and landscape aesthetics, human action is typically required for these services to actually flow to beneficiaries, e.g., produce being shipped to grocers; people venturing into nature to enjoy beautiful viewscapes. However, in the case of aesthetics, the data on which we based our mapping was informed by a viewshed analysis, which spatially quantified the potential for people to actually see areas all across the landscape, thereby incorporating an ES flow component. In the context of preventing or minimizing the impacts of flooding, waterflow regulation is provided (i.e., flows from supply to demand areas) when a supply area limits or delays the flow of water (Luck et al. 2009), which is typically a temporal dynamic dependent on seasonal temperature and/or weather patterns. Although the existing ES maps we use in this study are static spatial representations of potential supply areas, in the cases of waterflow regulation and landscape aesthetics, the distribution of potential spatial location of flows will still be captured by this mapping. Note also that ES flows are not equivalent to ES connectivity, the latter of which we are defining by the functional ecological interrelationships between different supply areas.

## Appendix 2 Original ES supply area mapping summary

Below we summarize the procedures for primary- and proxy-based mapping of the three ES used in our study, originally produced by Field et al. (2017). Ecosystem attributes were mapped, and their potential contribution to ES supply quantified, based on environmental functions that are known to underpin ES production; on explicit incorporation of perceived benefits to humans; or a combination of the two methods (Fig. 3; Jakeman and Letcher 2003, Vigerstol and Aukema 2011, Field et al. 2017). Spatial data sources for original maps included LULC indicators, remote sensing image interpretation, and were supported by some field-validations. For analytical consistency, raster data for original mapping were resampled to ∼29 m x 29 m resolution and assigned the identical spatial projection. Original maps were created using ArcMap 10.2 and 10.4 (ESRI 2011), and R (R Core Team 2013). For additional details see Field et al. (2017); data are available on the Open Science Framework (OSF; Field 2021).

Plant food agriculture (PA) is an economically and culturally important ES in our study area (e.g., Okanagan Valley Economic Development Society 2013, Kyle 2018), and its supply was mapped based on the spatial extent of all crop types used directly for human nutrition. These include tree fruits; vines and grapes; cereals and oilseeds; rotation crops; vegetables; berries; nut trees; and specialty foods, all of which are concentrated primarily in valley-bottom areas (Field et al. 2017). From these data, we dissolved boundaries between adjacent Agricultural Land Use Inventory (ALUI) polygons which, in some cases, resulted in different crop types being merged into a single node (MoAg 2017). We did this to generalize the mapping of connectivity between PA as a whole, and the other ES considered in this study, as the rationale for the mechanistic connections between PA and the other ES were consistent for all crop types. This outcome fits with our method of PA supply area valuation, which is based solely on potential crop area (ha) and not on crop type.

The terrestrial areas that provide waterflow regulation (WF) were mapped as a function of soil texture, slope, land use and land cover (LULC)-specific perviousness, and functionally relevant ecosystem types including floodplains, riparian areas, wetlands, and seasonally flooded fields (‘influential landscape features’ - ‘ILF’ herein). WF supply areas were defined as those that sustain water delivery in dedicated areas, and protect against flooding and draughts, both of which are persistent environmental concerns in the study region (Haughian et al. 2012). Several wetland areas were excluded from the Field et al. (2017) WF map due to the absence of soil texture data, which was one of the inputs for the waterflow infiltration model. Because wetlands are so critical to supporting WF and are relevant to connectivity mechanisms with several other ES, we added these areas back to our WF map by re-running the infiltration model under the assumption that all wetland areas without soils data have 100% saturated hydraulic conductivity, and then applying an ILF multiplier per Field et al. (2017). These resulting raster values ranged from 19 to 1200 (mean = 237.7; st.dev = 120.1), and wetlands coincident with mapped floodplains provided the highest-value WF supply areas in the region.

Lastly, landscape aesthetics (LA) supply areas were mapped based on models of perceived values of different LULC types in the region, on ‘visual condition’ ranging from preserved to manicured lands, and on the visibility of areas from various viewpoints across the case study region. LA supply areas spanned both terrestrial and large aquatic (i.e., lakes, rivers, manmade reservoirs) areas. We did not separate adjacent terrestrial from aquatic LA supply areas as 13 LULC (10 terrestrial; 3 aquatic) values were used as input for original LA mapping, in conjunction with two other valuation methods (i.e., tourism brochure assessment; viewshed analysis), and we aimed to keep supply area delineation methodologically as consistent as possible across different ES types (e.g., for amalgamation of immediately adjacent supply areas).

## Appendix 3 Major watersheds and sub-basins within and surrounding the case study landscape

Due to the large file size of the waterflow regulation (WF) data, supply area delineation steps were run separately for identified sub-basins (n = 118) in our case study area. To identify sub-basin catchment areas, we used BC Major Watershed, Fresh Water Atlas (FWA) Watersheds, and FWA Streams datasets (FLNRO 2017; see Field et al. 2017 for data source descriptions; datasets available at https://www.data.gov.bc.ca/). For major watersheds with significant (or complete) overlap with our study area, which would result in a large number of within-basin ES supply areas and therefore potentially lead to computational limitations, nested sub-basins were identified. These major watersheds included Kettle (west), Okanagan, Similkameen, and South Thompson rivers (Appendix B). FWA Watersheds with a common terminus into valley bottom waterbodies, verified using the FWA Streams dataset (FLNRO 2017), were merged. Several of the major watersheds (Columbia, Fraser, Kettle (east), Thompson, and Washington (Coast) rivers) overlapped with our study area primarily along its border; the overlapping portions of these watersheds were clipped and added to the sub-basin dataset (see Appendix B map). The high-value WF raster was then split by sub-basins using the nearest neighbour sampling technique and Split Raster tool in ArcMap. Each major watershed was assigned a unique ‘goal’ point location for LCP analyses, i.e., sub-basins within a major watershed shared the same goal point. In some cases, the mapped borders of the BC Major Watersheds and FWA Watersheds did not exhibit perfect overlap; therefore, following the high-value WF split exercise, the ArcGIS Erase tool was run by erasing the spatial extent of sub-basins from other BC Major Watersheds where they overlap. This ensured that WF supply areas were assigned to sub-basins by prioritizing the more detailed FWA Watershed dataset.

**Appendix 3 - Figure 1.**
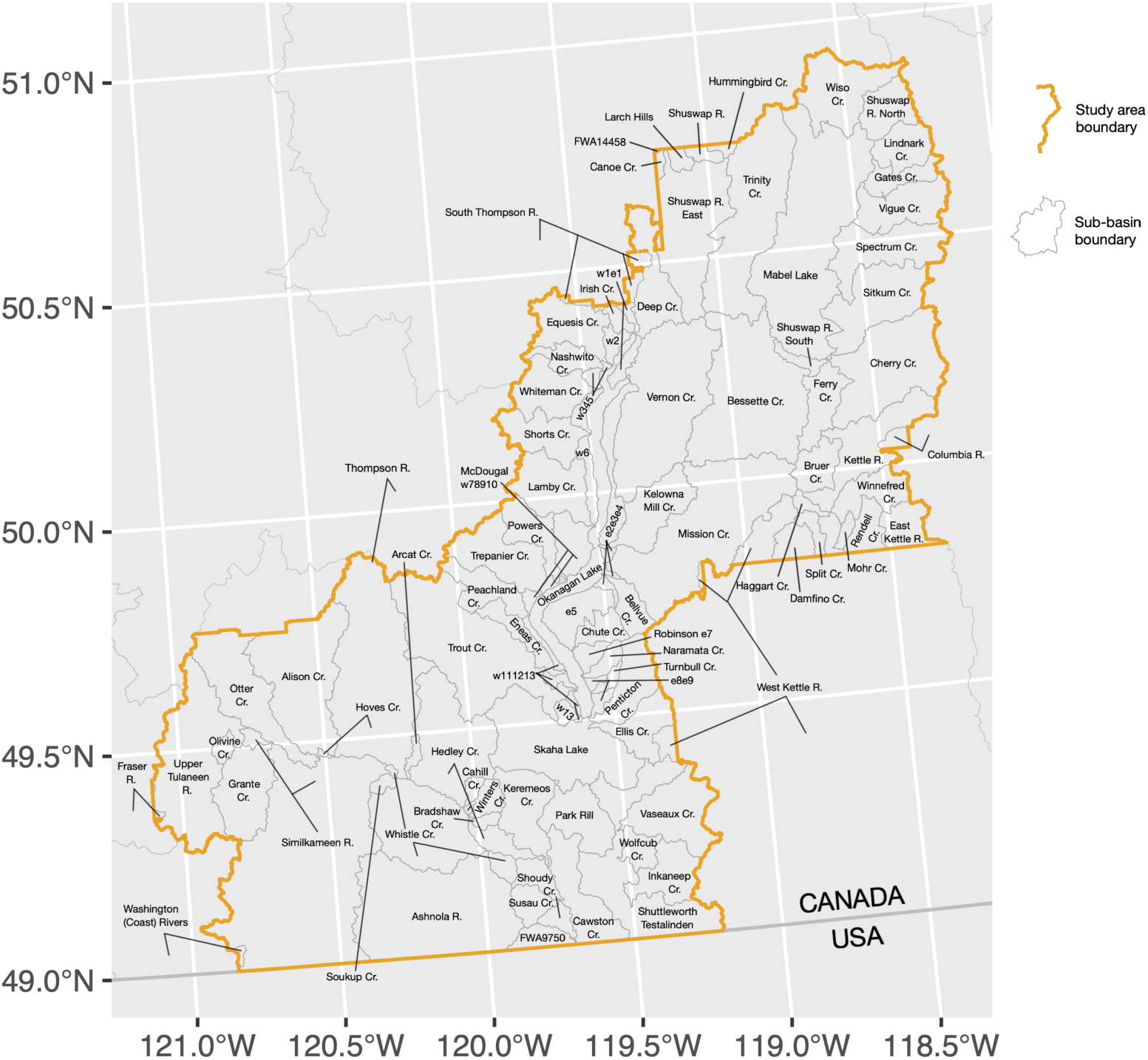
Map of major watersheds and sub-basins within and surrounding the case study landscape in southern interior British Columbia, Canada.

## Appendix 4 Detailed methods for building pairwise topographic corridors

Once initial LCPs were created, segments of LCP lines were erased where they were overlapped by a non-origin ES supply area polygon, and resulting disconnected lines were made into separate line features. Next, we deleted any lines that were deemed invalid from the perspective of real-world ES connectivity. Specifically, any resulting lines that intersected with the sub-basin goal point were deleted as we were only interested in retaining connections between pairs of ES supply areas. Certain ES polygons with an irregular shape had a centroid external to their polygon coverage, which resulted in line segments that initiated at the origin polygon centroid and terminated on the origin polygon border; these were also deleted as they did not represent links between a pair of ES supply areas. Irregular shaped nodes also sometimes yielded lines that were connected between two points on the parent-polygon border. These were retained to account for real-world feedbacks that may help maintain a supply area; however, we ensured that any such lines associated with two borders of an intersected polygon (i.e., where the line segment was part of a non-origin polygon) were deleted to avoid duplication with feedback links identified when separate analyses were run with the intersected polygon (in this example) as the origin (Fig. 6b).

For topographic corridor mapping we needed to address various rare analytical outcomes that became evident upon manual model validation. The LCP analyses resulted in some connections that violated landscape topography. For example, some LCPs from WF supply areas associated with relatively flat lands in the headwaters of one sub-basin (Bellevue Creek) were found to flow south to Okanagan Lake rather than flowing north as they would in reality (Appendix 2). This was due to the necessity of balancing the smallest possible raster resolution (20 m) with the overall large size of the study area for computational efficiency. As individual sub-basins were analyzed separately for topographic links, invalid linkages between sub-basins were not possible; however, we manually inspected all LCP results and removed any LCPs that violated downslope flow logic from subsequent analyses. Further, we did not incorporate certain rare spatial occurrences. These included instances of a smaller supply area inside bigger one; centroids captured by the incorrect buffer due to two or more centroids occurring close to one another; and LCP segments that resulted in feedback loops that occurred across non-origin supply areas (these loops were likely retained as feedback loops when such nodes served as LCP origins).

## Appendix 5 R script files (available on the OSF; Field 2021)

i. TOP-VALUE SUPPLY AREA NODE CREATION (file name: ‘nodes_waterflow_FORMAT.R’)
ii. OVERLAP LINK CREATION (file name: ‘links(overlap)_waterflow <> plantag_MASTER_FORMAT.R’)
iii. SUPPLY AREA CENTROID CREATION (PREP FOR LEAST COST PATH ’LCP’ ANALYSES) (file name: ‘links_waterflow_ALL_centroids_FORMAT.R’)
iv. CREATE LCP GOAL POINTS FOR EACH SUB-BASIN/MAJOR WATERSHED (file name: ‘links_waterflow_ALL_LCP goal pts_FORMAT.R’)
v. LEAST COST PATH ’LCP’ ANALYSIS FOR TOPOGRAPHIC LINK CREATION (file name: ‘links_waterflow_okanagan_LCPs_SUBBASIN_FORMAT.R’)
vi. TOPOGRAPHIC LINK CREATION (file name: ‘links_waterflow_okanagan_1_MASTER_FORMAT.R’)
vii. INFLUENTIAL LANDSCAPE FEATURES (ILF) LINKS (file name: ‘links_waterflow_okanagan_LCPs_ILF_FORMAT.R’)

## Appendix 6 Study limitations

We identified three primary limitations of our approach. Firstly, we only included three ES in our study and comparisons. Therefore, we are limited in the generalizations we can make, especially as they pertain to the specific locations, LULC- and ecosystem-relevance of the potential for connectivity ‘hotspots’. Investigating a limited number of ES is common among studies that model interactions among ES (Agudelo et al. 2020), with data limitations, complexity of socio-ecological process involved, and methodological gaps cited as barriers to inclusion of all ES (Kolosz et al. 2018). However, our choice to test only three ES was motivated by our goal to provide a straightforward case study of how each of the three broad ES categories (i.e., non ‘supporting’; MEA 2005) can be represented in the same study. Our approach is easily adaptable to including an unlimited number of ES, though the complexity in representing the functional connections between them will may increase disproportionately to the number of ES included, and limited data and/or gaps in our understanding of interaction mechanisms may preclude modelling of certain pairwise relationships (Field and Parrott 2017). Secondly, we only identified synergistic interactions among the case study ES we included, but no trade-offs were represented in our case study. The presence of potential trade-offs, as well as ecosystem dis-services (e.g., competition for water and pollination among different LULC types; spread of pests and diseases; Zhang et al. 2007), is of critical importance to informing management, as the optimization of all ES on a landscape is usually not simultaneously possible (e.g., Qiu and Turner 2013). We encourage future applications of our approach to represent trade-offs and negatively-valued functional connections between ES.

Lastly, we do not incorporate a measurement of uncertainty into our approach. For example, we did not attempt to directly assess spatial autocorrelation of our functional connectivity models with any other, potentially influential ecological processes, as we were interested in providing straight-forward and replicable rationale for mapping linkages; however, this precluded us from being able to parse the presence of shared drivers and potential artefacts in proxy or primary data. Additionally, the location and value of the identified connectivity corridors may be driven by the assumptions of original ES mapping and the threshold (top 50%) we used to delineate high-value supply areas. Several publications have suggested that incorporating uncertainty measures is necessary for producing reliable results to support decision making, and will lead to improved understanding of the system under study through identification of the most compelling findings (Seppelt et al. 2011, Hamel and Bryant 2017, Stritih et al. 2018). Sources of uncertainty considered in ES assessments are related to models of ecological processes; subjective choices of researchers and/or participants; and practical modelling skills and data quality (Gos and Lavorel 2012, Burkhard et al. 2013, Hou et al. 2018, Wang et al. 2018). To date, only a limited number of studies on ES interactions have incorporated measures of uncertainty and/or model validation (Boerema et al. 2017, Agudelo et al. 2020). As unconfirmed results are difficult to reliably assess, they are not as useful for direct practical applications (Agudelo et al. 2020). Studies with the express purpose of providing guidance for on-the-ground multi-ES planning should therefore incorporate metrics of uncertainty and model validation procedures.

## Appendix 7 Distribution of major LULC types across the case study region

**Appendix 7 - Figure 1.**
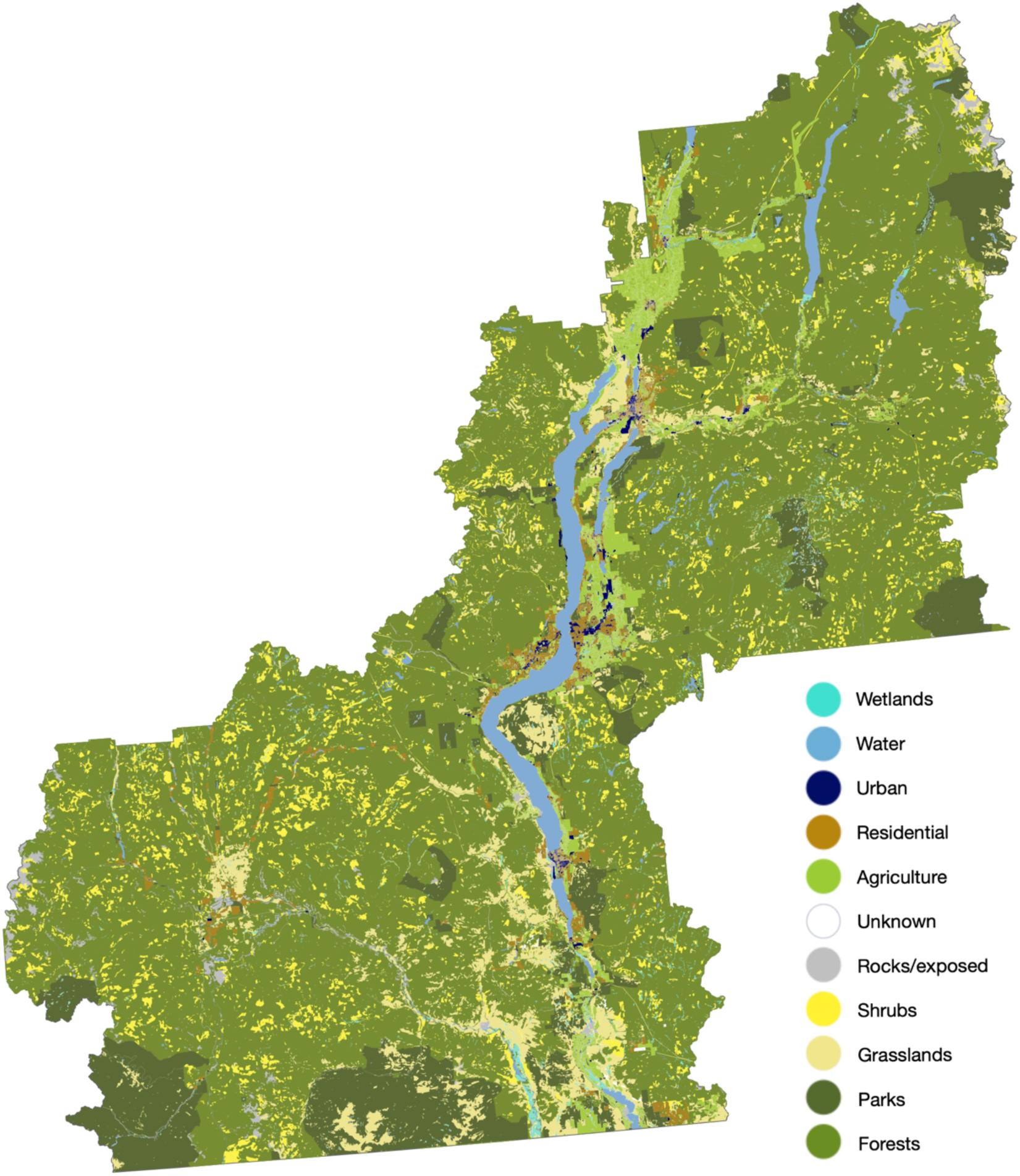
Map of the distribution of major LULC types across the case study region.

## Appendix 8 Summary tables of total areas and proportional coverages for major LULC types within each ES supply area and link type

**Appendix 8 - Table 1.** Percent of LULC overlapped by each ES supply area.

|  |  |  | High Value Supply Areas (Top 50%) |  |  |  |  |  |  |  |  |
| --- | --- | --- | --- | --- | --- | --- | --- | --- | --- | --- | --- |
|  |  | Study Area | PA |  |  | WF |  |  | LA |  |  |
| LULC Type | Total area (ha) | 2158001 | 12606 |  |  | 922425 |  |  | 1456241 |  |  |
|  |  | % of study area | Area (ha) | % of supply area | % of total LULC | Area (ha) | % of supply area | % of total LULC | Area (ha) | % of supply area | % of total LULC |
| Forest (incl. parks) | 1654215 | 76.65 | 30.85 | 0.24 | 1.86e-03 | 725852.30 | 78.69 | 43.88 | 1354434.00 | 93.01 | 81.88 |
| Park | 255964 | 11.86 | 4.25e-06 | 0 | 0 | 129588.70 | 14.05 | 50.63 | 190476.70 | 13.08 | 74.42 |
| Grassland | 147610 | 6.84 | 0.12 | 9.80e-04 | 8.37e-05 | 72050.87 | 7.81 | 48.81 | 3783.53 | 0.26 | 2.56 |
| Shrub | 124953 | 5.79 | 0 | 0 | 0 | 53425.76 | 5.79 | 42.76 | 3889.10 | 0.27 | 3.11 |
| Agriculture | 79769 | 3.70 | 12605.44 | 100.00 | 15.80 | 31338.00 | 3.40 | 39.29 | 241.00 | 0.02 | 0.30 |
| Water | 64272 | 2.98 | 0 | 0 | 0 | 0 | 0 | 0 | 63469.20 | 4.36 | 98.75 |
| Residential | 34897 | 1.62 | 3.08e-04 | 0 | 0 | 10990.55 | 1.19 | 31.49 | 241.44 | 0.02 | 0.69 |
| Rock / exposed | 27373 | 1.27 | 2.78e-04 | 0 | 0 | 5387.80 | 0.58 | 19.68 | 779.82 | 0.05 | 2.85 |
| Wetland | 18207 | 0.84 | 9.44e-06 | 0 | 0 | 18128.12 | 1.97 | 99.57 | 12586.06 | 0.86 | 69.13 |
| Urban | 6187 | 0.29 | 0 | 0 | 0 | 1918.28 | 0.21 | 31.01 | 27.75 | 1.91e-03 | 0.45 |
| Unknown | 1291 | 0.06 | 1.72 | 0.01 | 0.13 | 473.51 | 0.05 | 36.67 | 52.74 | 3.62e-03 | 4.08 |

**Appendix 8 - Table 2.** Percent of LULC overlapped by each topographic link.

|  |  | Topographic Links |  |  |  |  |  |  |  |  |  |
| --- | --- | --- | --- | --- | --- | --- | --- | --- | --- | --- | --- |
|  |  | Study Area | WF → PA |  |  | WF → WF |  |  | WF → LA |  |  |
| LULC Type | Total area (ha) | 2158001 | 4079 |  |  | 44449 |  |  | 4695 |  |  |
|  |  | % of study area | Area (ha) | % of link area | % of total LULC | Area (ha) | % of link area | % of total LULC | Area (ha) | % of link area | % of total LULC |
| Forest (incl. parks) | 1654215 | 76.65 | 2205.94 | 54.08 | 0.13 | 19069.77 | 42.90 | 1.15 | 2688.56 | 57.26 | 0.16 |
| Park | 255964 | 11.86 | 438.39 | 10.75 | 0.17 | 3146.14 | 7.08 | 1.23 | 682.44 | 14.54 | 0.27 |
| Grassland | 147610 | 6.84 | 433.05 | 10.62 | 0.29 | 3065.90 | 6.90 | 2.08 | 477.85 | 10.18 | 0.32 |
| Shrub | 124953 | 5.79 | 102.67 | 2.52 | 0.08 | 1230.22 | 2.77 | 0.98 | 195.80 | 4.17 | 0.16 |
| Agriculture | 79769 | 3.70 | 807.02 | 19.79 | 1.01 | 2329.46 | 5.24 | 2.92 | 596.03 | 12.70 | 0.75 |
| Water | 64272 | 2.98 | 127.65 | 3.13 | 0.20 | 15664.49 | 35.24 | 24.37 | 184.83 | 3.94 | 0.29 |
| Residential | 34897 | 1.62 | 218.51 | 5.36 | 0.63 | 1368.65 | 3.08 | 3.92 | 304.98 | 6.50 | 0.87 |
| Rock / exposed | 27373 | 1.27 | 46.70 | 1.14 | 0.17 | 671.99 | 1.51 | 2.45 | 112.70 | 2.40 | 0.41 |
| Wetland | 18207 | 0.84 | 56.51 | 1.39 | 0.31 | 732.48 | 1.65 | 4.02 | 51.79 | 1.10 | 0.28 |
| Urban | 6187 | 0.29 | 43.92 | 1.08 | 0.71 | 185.53 | 0.42 | 3.00 | 45.69 | 0.97 | 0.74 |
| Unknown | 1291 | 0.06 | 1.07 | 0.03 | 0.08 | 2.83 | 0.01 | 0.22 | 1.72 | 0.04 | 0.13 |

**Appendix 8 - Table 3.**
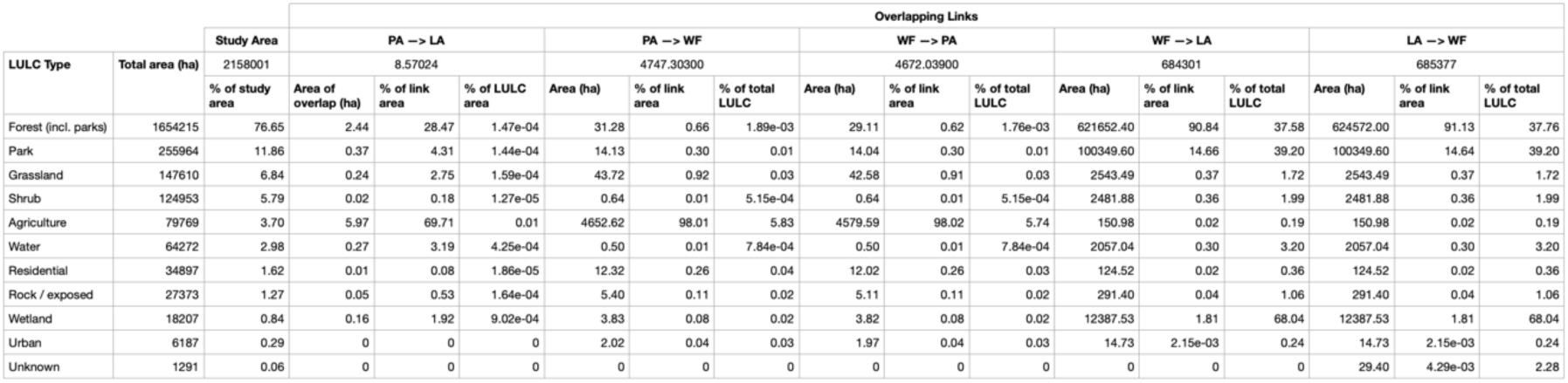
Percent of LULC overlapped by each overlapping links.

## References

Agudelo C. A. R., S. L. H. Bustos, and C. A. P. Moreno. 2020. Modeling interactions among multiple ecosystem services. A critical review. Ecological Modelling 429:109103. https://doi.org/10.1016/j.ecolmodel.2020.109103

Alemu J. B., D. R. Richards, L. Y. Gaw, M. Masoudi, Y. Nathan, and D. A. Friess. 2021. Identifying spatial patterns and interactions among multiple ecosystem services in an urban mangrove landscape. Ecological Indicators 121:107042. https://doi.org/10.1016/j.ecolind.2020.107042

Alexander R. B., E. W. Boyer, R. A. Smith, G. E. Schwarz, and R. B. Moore. 2007. The role of headwater streams in downstream water quality. JAWRA Journal of the American Water Resources Association 43(1):41–59. https://doi.org/10.1111/j.1752-1688.2007.00005.x

Anderson B. J., P. R. Armsworth, F. Eigenbrod, C. D. Thomas, S. Gillings, A. Heinemeyer, D. B. Roy, and K. J. Gaston. 2009. Spatial covariance between biodiversity and other ecosystem service priorities. Journal of Applied Ecology 46(4):888–896. https://doi.org/10.1111/j.1365-2664.2009.01666.x

Bagstad K. J., D. J. Semmens, S. Waage, and R. Winthrop. 2013. A comparative assessment of decision-support tools for ecosystem services quantification and valuation. Ecosystem services 5:27–39. https://doi.org/10.1016/j.ecoser.2013.07.004

Bangash R. F., A. Passuello, M. Sanchez-Canales, M. Terrado, A. López, F. J. Elorza, G. Ziv, V. Acuña, and M. Schuhmacher. 2013. Ecosystem services in Mediterranean river basin: climate change impact on water provisioning and erosion control. Science of the Total Environment 458:246–255. https://doi.org/10.1016/j.scitotenv.2013.04.025

Batary P., A. Baldi, M. Sarospataki, F. Kohler, J. Verhulst, E. Knop, F. Herzog, and D. Kleijn. 2010. Effect of conservation management on bees and insect-pollinated grassland plant communities in three European countries. Agriculture, Ecosystems & Environment 136(1):35–39. https://doi.org/10.1016/j.agee.2009.11.004

Beier P., D. R. Majka, and S. L. Newell. 2009. Uncertainty analysis of least-cost modeling for designing wildlife linkages. Ecological Applications 19(8):2067–2077. https://doi.org/10.1890/08-1898.1

Bennett E. M., G. D. Peterson, and L. J. Gordon. 2009. Understanding relationships among multiple ecosystem services. Ecology Letters 12(12):1394–1404. https://doi.org/10.1111/j.1461-0248.2009.01387.x

Bennett E. M., J. Baird, H. Baulch, R. Chaplin-Kramer, E. Fraser, P. Loring, P. Morrison, L. Parrott, K. Sherren, and K. J. Winkler. 2021. Ecosystem services and the resilience of agricultural landscapes. Pages 1–43 *In* Advances in Ecological Research, Elsevier. https://doi.org/10.1016/bs.aecr.2021.01.001

Berdugo M., M. Delgado-Baquerizo, S. Soliveres, R. Hernández-Clemente, Y. Zhao, J. J. Gaitán, N. Gross, H. Saiz, V. Maire, and A. Lehman. 2020. Global ecosystem thresholds driven by aridity. Science 367(6479):787–790. https://doi.org/10.1126/science.aay5958

Beza B. B. 2010. The aesthetic value of a mountain landscape: A study of the Mt. Everest Trek. Landscape and Urban Planning 97(4):306–317. https://doi.org/10.1016/j.landurbplan.2010.07.003

Bialonski S., M. Horstmann, and K. Lehnertz. 2010. From brain to earth and climate systems: Small-world interaction networks or not? Chaos: An Interdisciplinary Journal of Nonlinear Science 20(1):013134. https://doi.org/10.1063/1.3360561

Biesmeijer J. C., S. P. Roberts, M. Reemer, R. Ohlemuller, M. Edwards, T. Peeters, A. P. Schaffers, S. G. Potts, R. Kleukers, C. D. Thomas, J. Settele, and W. E. Kunin. 2006. Parallel declines in pollinators and insect-pollinated plants in Britain and the Netherlands. Science 313(5785):351–354. https://doi.org/10.1126/science.1127863

Biggs R., M. Schlüter, D. Biggs, E. L. Bohensky, S. BurnSilver, G. Cundill, V. Dakos, T. M. Daw, L. S. Evans, and K. Kotschy. 2012. Toward principles for enhancing the resilience of ecosystem services. Annual Review of Environment and Resources 37:421–448. https://doi.org/10.1146/annurev-environ-051211-123836

Birkhofer K., G. K. Andersson, J. Bengtsson, R. Bommarco, J. Dänhardt, B. Ekbom, J. Ekroos, T. Hahn, K. Hedlund, and A. M. Jönsson. 2018. Relationships between multiple biodiversity components and ecosystem services along a landscape complexity gradient. Biological Conservation 218:247–253. https://doi.org/10.1016/j.biocon.2017.12.027

Bivand R., C. Rundel, E. Pebesma, R. Stuetz, K. O. Hufthammer, and M. R. Bivand. 2017. Package ‘rgeos’. The Comprehensive R Archive Network (CRAN). https://r-forge.r-project.org/projects/rgeos/

Bivand R., T. Keitt, B. Rowlingson, E. Pebesma, M. Sumner, R. Hijmans, E. Rouault, and M. R. Bivand. 2015. Package ‘rgdal’. Bindings for the Geospatial Data Abstraction Library.https://cran.r-project.org/web/packages/rgdal/index.html

Bivand R. S., E. J. Pebesma, V. Gomez-Rubio, and E. J. Pebesma. 2013. Applied spatial data analysis with R, Volume 2. Springer, New York. https://doi.org/10.1007/978-1-4614-7618-4

Blackstock M. 2001. Water: A First Nations’ Spiritual and Ecological Perspective. BC Journal of Ecosystems and Management 1(1):1–14.

Boesing A. L., P. R. Prist, J. Barreto, C. Hohlenwerger, M. Maron, J. R. Rhodes, E. Romanini, L. R. Tambosi, M. Vidal, and J. P. Metzger. 2020. Ecosystem services at risk: integrating spatiotemporal dynamics of supply and demand to promote long-term provision. One Earth 3(6):704–713. https://doi.org/10.1016/j.oneear.2020.11.003

Bohan D. A., A. Raybould, C. Mulder, G. Woodward, A. Tamaddoni-Nezhad, N. Bluthgen, M. J. Pocock, S. Muggleton, D. M. Evans, and J. Astegiano. 2013. Networking agroecology: integrating the diversity of agroecosystem interactions. Advances in Ecological Research 49:1–67. https://doi.org/10.1016/B978-0-12-420002-9.00001-9

Boyd J., S. Banzhaf. 2007. What are ecosystem services? The need for standardized environmental accounting units. Ecological Economics 63(2):616–626. https://doi.org/10.1016/j.ecolecon.2007.01.002

Brander L. M., R. J. Florax, and J. E. Vermaat. 2006. The empirics of wetland valuation: a comprehensive summary and a meta-analysis of the literature. Environmental and Resource Economics 33(2):223–250. https://doi.org/10.1007/s10640-005-3104-4

Brauman K. A., G. C. Daily, T. K. Duarte, and H. A. Mooney. 2007. The nature and value of ecosystem services: an overview highlighting hydrologic services. Annu.Rev.Environ.Resour. 32:67–98. https://doi.org/10.1146/annurev.energy.32.031306.102758

Brooks C. 2003. A scalar analysis of landscape connectivity. Oikos :433–439.

Burmil S., T. C. Daniel, and J. D. Hetherington. 1999. Human values and perceptions of water in arid landscapes. Landscape and Urban Planning 44(2):99–109. https://doi.org/10.1016/S0169-2046(99)00007-9

Butchart S. H., M. Walpole, B. Collen, A. van Strien, J. P. Scharlemann, R. E. Almond, J. E. Baillie, B. Bomhard, C. Brown, J. Bruno, K. E. Carpenter, G. M. Carr, J. Chanson, A. M. Chenery, J. Csirke, N. C. Davidson, F. Dentener, M. Foster, A. Galli, J. N. Galloway, P. Genovesi, R. D. Gregory, M. Hockings, V. Kapos, J. F. Lamarque, F. Leverington, J. Loh, M. A. McGeoch, L. McRae, A. Minasyan, M. Hernandez Morcillo, T. E. Oldfield, D. Pauly, S. Quader, C. Revenga, J. R. Sauer, B. Skolnik, D. Spear, D. Stanwell-Smith, S. N. Stuart, A. Symes, M. Tierney, T. D. Tyrrell, J. C. Vie, and R. Watson. 2010. Global biodiversity: indicators of recent declines. Science 328(5982):1164–1168. https://doi.org/10.1126/science.1187512

Cadotte M. W., K. Carscadden, and N. Mirotchnick. 2011. Beyond species: functional diversity and the maintenance of ecological processes and services. Journal of Applied Ecology 48(5):1079–1087. https://doi.org/10.1111/j.1365-2664.2011.02048.x

Carpenter S. R., E. G. Booth, S. Gillon, C. J. Kucharik, S. Loheide, A. S. Mase, M. Motew, J. Qiu, A. R. Rissman, and J. Seifert. 2015. Plausible futures of a social-ecological system: Yahara watershed, Wisconsin, USA. Ecology and Society 20(2):10. http://dx.doi.org/10.5751/ES-07433-200210

Carpenter S. R., H. A. Mooney, J. Agard, D. Capistrano, R. S. Defries, S. Diaz, T. Dietz, A. K. Duraiappah, A. Oteng-Yeboah, H. M. Pereira, C. Perrings, W. V. Reid, J. Sarukhan, R. J. Scholes, and A. Whyte. 2009. Science for managing ecosystem services: Beyond the Millennium Ecosystem Assessment. Proceedings of the National Academy of Sciences of the United States of America 106(5):1305–1312. https://doi.org/10.1073/pnas.0808772106

Caslys Consulting Ltd. A biodiversity conservation analysis summary for the Okanagan region. 2013. Okanagan Collaborative Conservation Program and South Okanagan Similkameen Conservation Program, Kelowna, British Columbia.

Chi Y., H. Shi, W. Zheng, J. Sun, and Z. Fu. 2018. Spatiotemporal characteristics and ecological effects of the human interference index of the Yellow River Delta in the last 30 years. Ecological Indicators 89:880–892. https://doi.org/10.1016/j.ecolind.2017.12.025

Cord A. F., N. Schwarz, R. Seppelt, M. Volk, and M. Schröter. 2019. Introduction to Part III: Trade-Offs and Synergies Among Ecosystem Services. Pages 245–249 *In* M. Schröter, A. Bonn, S. Klotz, R. Seppelt, C. Baessler, editors. Atlas of Ecosystem Services, Springer, Cham. https://doi.org/10.1007/978-3-319-96229-0_38

Cressie N. 1993. Statistics for spatial data. John Wiley & Sons, Inc., New York.

Crossman N. D., B. Burkhard, S. Nedkov, L. Willemen, K. Petz, I. Palomo, E. G. Drakou, B. Martín-Lopez, T. McPhearson, and K. Boyanova. 2013. A blueprint for mapping and modelling ecosystem services. Ecosystem Services 4:4–14. https://doi.org/10.1016/j.ecoser.2013.02.001

Cui B., Z. Zhang, and X. Lei. 2012. Implementation of diversified ecological networks to strengthen wetland conservation. Clean–Soil, Air, Water 40(10):1015–1026. https://doi.org/10.1002/clen.201200026

Cumming G. S., D. H. Cumming, and C. L. Redman. 2006. Scale mismatches in social-ecological systems: causes, consequences, and solutions. Ecology and society 11(1).

Dabney S. M. 1998. Cover crop impacts on watershed hydrology. Journal of Soil and Water Conservation 53(3):207–213.

Daily G. C., S. Polasky, J. Goldstein, P. M. Kareiva, H. A. Mooney, L. Pejchar, T. H. Ricketts, J. Salzman, and R. Shallenberger. 2009. Ecosystem services in decision making: time to deliver. Frontiers in Ecology and the Environment 7(1):21–28. https://doi.org/10.1890/080025

Dalgaard T., N. J. Hutchings, and J. R. Porter. 2003. Agroecology, scaling and interdisciplinarity. Agriculture, Ecosystems & Environment 100(1):39–51. https://doi.org/10.1016/S0167-8809(03)00152-X

Daniel T. C., A. Muhar, A. Arnberger, O. Aznar, J. W. Boyd, K. M. Chan, R. Costanza, T. Elmqvist, C. G. Flint, P. H. Gobster, A. Gret-Regamey, R. Lave, S. Muhar, M. Penker, R. G. Ribe, T. Schauppenlehner, T. Sikor, I. Soloviy, M. Spierenburg, K. Taczanowska, J. Tam, and A. von der Dunk. 2012. Contributions of cultural services to the ecosystem services agenda. Proceedings of the National Academy of Sciences of the United States of America 109(23):8812–8819. https://doi.org/10.1073/pnas.1114773109

De Groot R. S., R. Alkemade, L. Braat, L. Hein, and L. Willemen. 2010. Challenges in integrating the concept of ecosystem services and values in landscape planning, management and decision making. Ecological complexity 7(3):260–272. https://doi.org/10.1016/j.ecocom.2009.10.006

De Loë R., R. Kreutzwiser, and L. Moraru. 2001. Adaptation options for the near term: climate change and the Canadian water sector. Global Environmental Change 11(3):231–245. https://doi.org/10.1016/S0959-3780(00)00053-4

Debinski D. M., R. D. Holt. 2000. A survey and overview of habitat fragmentation experiments. Conservation Biology 14(2):342–355. https://doi.org/10.1046/j.1523-1739.2000.98081.x

Dee L. E., S. Allesina, A. Bonn, A. Eklöf, S. D. Gaines, J. Hines, U. Jacob, E. McDonald-Madden, H. Possingham, and M. Schröter. 2017. Operationalizing network theory for ecosystem service assessments. Trends in ecology & evolution 32(2):118–130. https://doi.org/10.1016/j.tree.2016.10.011

DeLaney T. A. 1995. Benefits to downstream flood attenuation and water quality as a result of constructed wetlands in agricultural landscapes. Journal of Soil and Water Conservation 50(6):620–626.

Dobbs C., D. Kendal, and C.R. Nitschke. 2014. Multiple ecosystem services and disservices of the urban forest establishing their connections with landscape structure and sociodemographics. Ecological Indicators. 43:44–55. https://doi.org/10.1016/j.ecolind.2014.02.007

Dodds W. K., R. M. Oakes. 2008. Headwater influences on downstream water quality. Environmental Management 41(3):367–377. https://doi.org/10.1007/s00267-007-9033-y

D’Odorico P., G. S. Okin, and B. T. Bestelmeyer. 2012. A synthetic review of feedbacks and drivers of shrub encroachment in arid grasslands. Ecohydrology 5(5):520–530. https://doi.org/10.1002/eco.259

Dutta S., M. Sahana, and S. K. Guchhait. 2017. Assessing anthropogenic disturbance on forest health based on fragment grading in Durgapur Forest Range, West Bengal, India. Spatial Information Research 25(3):501–512. https://doi.org/10.1007/s41324-017-0117-3

Egoh B., B. Reyers, M. Rouget, D. M. Richardson, D. C. Le Maitre, and A. S. van Jaarsveld. 2008. Mapping ecosystem services for planning and management. Agriculture, Ecosystems & Environment 127(1):135–140. https://doi.org/10.1016/j.agee.2008.03.013

Ekroos J., O. Olsson, M. Rundlöf, F. Wätzold, and H. G. Smith. 2014. Optimizing agri-environment schemes for biodiversity, ecosystem services or both? Biological Conservation 172:65–71. https://doi.org/10.1016/j.biocon.2014.02.013

Ernstson H., S. Barthel, E. Andersson, and S. T. Borgström. 2010. Scale-crossing brokers and network governance of urban ecosystem services: the case of Stockholm. Ecology and Society 15(4):28.

ESRI. 2011. ArcGIS Desktop. Redlands, CA: Environmental Systems Research Institute. Release 10.

Fahrig L. 2003. Effects of habitat fragmentation on biodiversity. Annual Review of Ecology, Evolution, and Systematics 34:487–515. https://doi.org/10.1146/annurev.ecolsys.34.011802.132419

Fall A., M. Fortin, M. Manseau, and D. O’Brien. 2007. Spatial graphs: principles and applications for habitat connectivity. Ecosystems 10(3):448–461. https://doi.org/10.1007/s10021-007-9038-7

Field R. D. 2021. Mapping the functional connectivity of ecosystem services supply across a regional landscape - Data and scripts. Open Science Framework https://doi.org/10.17605/OSF.IO/9S4RM

Field R, Kyle C, Pedersen S, Parrott L. Ecosystem Services Mapping to Support Environmental Conservation Planning Decisions in the Okanagan Region: Year 1 Progress Report. Kelowna, BC: Prepared by: Okanagan Institute for Biodiversity, Resilience and Ecosystem Services (BRASES), Prepared for: Environment and Climate Change Canada.; 2017. Report nr GCXE17P178. 39 p.

Field R. D., L. Parrott. 2017. Multi-ecosystem services networks: A new perspective for assessing landscape connectivity and resilience. Ecological Complexity 32:31–41. https://doi.org/10.1016/j.ecocom.2017.08.004

Fischer J., D. B. Lindenmayer, and A. D. Manning. 2006. Biodiversity, ecosystem function, and resilience: ten guiding principles for commodity production landscapes. Frontiers in Ecology and the Environment 4(2):80–86. https://doi.org/10.1890/1540-9295(2006)004[0080:BEFART]2.0.CO;2

Fleischner T. L. 1994. Ecological costs of livestock grazing in western North America. Conservation Biology 8(3):629–644. https://doi.org/10.1046/j.1523-1739.1994.08030629.x

FLNRO (BC Ministry of Forests, Lands and Natural Resource Operations). 2017. Freshwater Atlas. BC Data Catalogue. https://data.gov.bc.ca/

Foley J. A., N. Ramankutty, K. A. Brauman, E. S. Cassidy, J. S. Gerber, M. Johnston, N. D. Mueller, C. O’Connell, D. K. Ray, and P. C. West. 2011. Solutions for a cultivated planet. Nature 478(7369):337–342. https://doi.org/10.1038/nature10452

Foley J. A., R. Defries, G. P. Asner, C. Barford, G. Bonan, S. R. Carpenter, F. S. Chapin, M. T. Coe, G. C. Daily, H. K. Gibbs, J. H. Helkowski, T. Holloway, E. A. Howard, C. J. Kucharik, C. Monfreda, J. A. Patz, I. C. Prentice, N. Ramankutty, and P. K. Snyder. 2005. Global consequences of land use. Science 309(5734):570–574. https://doi.org/10.1126/science.1111772

Fortuna M. A., C. Gomez-Rodriguez, and J. Bascompte. 2006. Spatial network structure and amphibian persistence in stochastic environments. Proceedings of the Royal Society B: Biological Sciences 273(1592):1429–1434. https://doi.org/10.1098/rspb.2005.3448

Freeman M. C., C. M. Pringle, and C. R. Jackson. 2007. Hydrologic connectivity and the contribution of stream headwaters to ecological integrity at regional scales. JAWRA Journal of the American Water Resources Association 43(1):5–14. https://doi.org/10.1111/j.1752-1688.2007.00002.x

Friess D. A. 2016. Ecosystem services and disservices of mangrove forests: insights from historical colonial observations. Forests 7(9):183. https://doi.org/10.3390/f7090183

Fu B., S. Wang, C. Su, and M. Forsius. 2013. Linking ecosystem processes and ecosystem services. Current Opinion in Environmental Sustainability 5(1):4–10. https://doi.org/10.1016/j.cosust.2012.12.002

Fu B., J. Wang, L. Chen, and Y. Qiu. 2003. The effects of land use on soil moisture variation in the Danangou catchment of the Loess Plateau, China. Catena 54(1-2):197–213. https://doi.org/10.1016/S0341-8162(03)00065-1

Fu B., Y. Wang, Y. Lu, C. He, L. Chen, and C. Song. 2009. The effects of land-use combinations on soil erosion: a case study in the Loess Plateau of China. Progress in Physical Geography 33(6):793–804. https://doi.org/10.1177/0309133309350264

Gardner R. H. 1999. RULE: map generation and a spatial analysis program. Pages 280–303 *In* J. M. Klopatek and R. H. Gardner, editors. Landscape Ecological Analysis, Springer-Verlag, New York.

Gilarranz L. J., J. M. Pastor, and J. Galeano. 2012. The architecture of weighted mutualistic networks. Oikos 121(7):1154–1162. https://doi.org/10.1111/j.1600-0706.2011.19592.x

Gonzalez A., N. Mouquet, and M. Loreau. 2009. Biodiversity as spatial insurance: the effects of habitat fragmentation and dispersal on ecosystem functioning. Pages 134–146 *In* S. Naeem, D.E. Bunker, A. Hector, M. Loreau, C. Perrings, editors. Biodiversity, ecosystem functioning and ecosystem services, Oxford University Press, New York.

Grêt-Regamey A., E. Sirén, S. H. Brunner, and B. Weibel. 2017. Review of decision support tools to operationalize the ecosystem services concept. Ecosystem Services 26:306–315. https://doi.org/10.1016/j.ecoser.2016.10.012

Guerry A. D., S. Polasky, J. Lubchenco, R. Chaplin-Kramer, G. C. Daily, R. Griffin, M. Ruckelshaus, I. J. Bateman, A. Duraiappah, T. Elmqvist, M. W. Feldman, C. Folke, J. Hoekstra, P. M. Kareiva, B. L. Keeler, S. Li, E. McKenzie, Z. Ouyang, B. Reyers, T. H. Ricketts, J. Rockstrom, H. Tallis, and B. Vira. 2015. Natural capital and ecosystem services informing decisions: From promise to practice. Proceedings of the National Academy of Sciences of the United States of America 112(24):7348–7355. https://doi.org/10.1073/pnas.1503751112

Haase D., N. Schwarz, M. Strohbach, F. Kroll, and R. Seppelt. 2012. Synergies, trade-offs, and losses of ecosystem services in urban regions: an integrated multiscale framework applied to the Leipzig-Halle Region, Germany. Ecology and Society 17(3):22.

Heckmann T., W. Schwanghart. 2013. Geomorphic coupling and sediment connectivity in an alpine catchment—Exploring sediment cascades using graph theory. Geomorphology 182:89–103. https://doi.org/10.1016/j.geomorph.2012.10.033

Heckmann T., W. Schwanghart, and J. D. Phillips. 2015. Graph theory—Recent developments of its application in geomorphology. Geomorphology 243:130–146. https://doi.org/10.1016/j.geomorph.2014.12.024

Hijmans R. J., J. van Etten. 2012. raster: Geographic analysis and modeling with raster data. R package version 2.0-12. http://CRAN.R-project.org/package=raster.

Hines J., van der Putten, Wim H, G. B. De Deyn, C. Wagg, W. Voigt, C. Mulder, W. W. Weisser, J. Engel, C. Melian, and S. Scheu. 2015. Towards an integration of biodiversity–ecosystem functioning and food web theory to evaluate relationships between multiple ecosystem services. Advances in Ecological Research 53:161–199. https://doi.org/10.1016/bs.aecr.2015.09.001

Horvát, E-A, K.A. Zweig 2012. One-mode projections of multiplex bipartite graphs. Pages 599–606 *In* Proceedings of the 2012 International Conference on Advances in Social Networks Analysis and Mining (ASONAM 2012), August 26-29, IEEE Computer Society, Washington, DC. https://doi.org/10.1109/ASONAM.2012.101

Houlahan J. E., C. S. Findlay. 2004. Estimating the ‘critical’ distance at which adjacent land-use degrades wetland water and sediment quality. Landscape Ecology 19(6):677–690. https://doi.org/10.1023/B:LAND.0000042912.87067.35

Ingram J., P. Gregory, and A. Izac. 2008. The role of agronomic research in climate change and food security policy. Agriculture, Ecosystems & Environment 126(1-2):4–12. https://doi.org/10.1016/j.agee.2008.01.009

Jackson R. B., S. R. Carpenter, C. N. Dahm, D. M. McKnight, R. J. Naiman, S. L. Postel, and S. W. Running. 2001. Water in a changing world. Ecological Applications 11(4):1027–1045. https://doi.org/10.1890/1051-0761(2001)011[1027:WIACW]2.0.CO;2

Janssen M. A., Ö. Bodin, J. M. Anderies, T. Elmqvist, H. Ernstson, R. R. McAllister, P. Olsson, and P. Ryan. 2006. Toward a network perspective of the study of resilience in social-ecological systems. Ecology and Society 11(1):15. http://www.jstor.org/stable/26267803

Kerr J. T., J. Cihlar. 2004. Patterns and causes of species endangerment in Canada. Ecological Applications 14(3):743–753. https://doi.org/10.1890/02-5117

Kolosz B., I. Athanasiadis, G. Cadisch, T. Dawson, C. Giupponi, M. Honzák, J. Martinez-Lopez, A. Marvuglia, V. Mojtahed, and K. B. Ogutu. 2018. Conceptual advancement of socio-ecological modelling of ecosystem services for re-evaluating Brownfield land. Ecosystem Services 33:29–39. https://doi.org/10.1016/j.ecoser.2018.08.003

Kort J. 1988. 9. Benefits of windbreaks to field and forage crops. Agriculture, Ecosystems & Environment 22:165–190. https://doi.org/10.1016/0167-8809(88)90017-5

Kraus E. 2014. Gloria Pungetti, Gonzalo Oviedo, and Della Hooke (eds.), Sacred species and sites: advances in biocultural conservation. Landscape Ecology 29(8):1461–1462. https://doi.org/10.1007/s10980-014-0072-5

Kremen C., R. S. Ostfeld. 2005. A call to ecologists: measuring, analyzing, and managing ecosystem services. Frontiers in Ecology and the Environment 3(10):540–548. https://doi.org/10.1890/1540-9295(2005)003[0540:ACTEMA]2.0.CO;2

Kremen C., N. M. Williams, M. A. Aizen, B. Gemmill-Herren, G. LeBuhn, R. Minckley, L. Packer, S. G. Potts, T. Roulston, and I. Steffan-Dewenter. 2007. Pollination and other ecosystem services produced by mobile organisms: a conceptual framework for the effects of land-use change. Ecology Letters 10(4):299–314. https://doi.org/10.1111/j.1461-0248.2007.01018.x

Kyle C. 2018. Chinese and Japanese Market Gardening in the North and Central Okanagan Valley, British Columbia. BC Studies: The British Columbian Quarterly 200:241–272.

Leibold M. A., M. Holyoak, N. Mouquet, P. Amarasekare, J. M. Chase, M. F. Hoopes, R. D. Holt, J. B. Shurin, R. Law, and D. Tilman. 2004. The metacommunity concept: a framework for multi-scale community ecology. Ecology Letters 7(7):601–613. https://doi.org/10.1111/j.1461-0248.2004.00608.x

Lewin-Koh N. J., R. Bivand, J. Pebesma, E. Archer, A. Baddeley, D. G. Giraudoux, V. G. Rubio, P. Hausmann, K. O. Hufthammer, and T. Jagger. 2012. Package ‘maptools’. http://cran.r-project.org/web/packages/maptools/maptools.pdf

Li Y., L. Zhang, J. Qiu, J. Yan, L. Wan, P. Wang, N. Hu, W. Cheng, and B. Fu. 2017. Spatially explicit quantification of the interactions among ecosystem services. Landscape Ecology 32(6):1181–1199. https://doi.org/10.1007/s10980-017-0527-6

Liu S., D. Wang, H. Li, W. Li, and Q. Wang. 2017. Ecological land fragmentation evaluation and dynamic change of a typical black soil farming area in northeast China. Sustainability 9(2):300. https://doi.org/10.3390/su9020300

Lonsdorf E., T. Ricketts, C. Kremen, R. Winfree, S. Greenleaf, and N. Williams. 2011. Crop pollination services. Pages 168–187 *In* P. Karieva, H. Tallis, T. H. Ricketts, G. C. Daily, and S. Polasky, editors. Natural Capital: Theory and practice of mapping ecosystem services, Oxford University Press, Oxford, UK.

Loomis J. 2002. Quantifying recreation use values from removing dams and restoring free-flowing rivers: A contingent behavior travel cost demand model for the Lower Snake River. Water Resources Research 38(6):2–1-2-8. https://doi.org/10.1029/2000WR000136

Loos J., P. Batáry, I. Grass, C. Westphal, S. Bänsch, A. B. Baillod, A. L. Hass, J. Rosa, and T. Tscharntke. 2019. Vulnerability of ecosystem services in farmland depends on landscape management. Pages 91–96 *In* M. Schröter, A. Bonn, S. Klotz, R. Seppelt, C. Baessler, editors. Atlas of Ecosystem Services, Springer, Cham. https://doi.org/10.1007/978-3-319-96229-0_15

Lorilla R. S., K. Poirazidis, S. Kalogirou, V. Detsis, and A. Martinis. 2018. Assessment of the spatial dynamics and interactions among multiple ecosystem services to promote effective policy making across mediterranean island landscapes. Sustainability 10(9):3285. https://doi.org/10.3390/su10093285

Lyons K. G., C. Brigham, B. Traut, and M. W. Schwartz. 2005. Rare species and ecosystem functioning. Conservation Biology 19(4):1019–1024. https://doi.org/10.1111/j.1523-1739.2005.00106.x

Maestre F. T., D. J. Eldridge, S. Soliveres, S. Kéfi, M. Delgado-Baquerizo, M. A. Bowker, P. García-Palacios, J. Gaitán, A. Gallardo, and R. Lázaro. 2016. Structure and functioning of dryland ecosystems in a changing world. Annual review of ecology, evolution, and systematics 47:215–237. https://doi.org/10.1146/annurev-ecolsys-121415-032311

Mäler K., S. Aniyar, and Å. Jansson. 2013. Accounting for the value of ecosystem assets and their services. Pages 187–205 *In* R. M. Hassen and E. D. Mungatana, editors. Implementing Environmental Accounts: Case Studies from Eastern and Southern Africa, Springer, Dordrecht. https://doi.org/10.1007/978-94-007-5323-5_7

Margosian M. L., K. A. Garrett, J. S. Hutchinson, and K. A. With. 2009. Connectivity of the American agricultural landscape: assessing the national risk of crop pest and disease spread. Bioscience 59(2):141–151. https://doi.org/10.1525/bio.2009.59.2.7

Marini L., M. Quaranta, P. Fontana, J. C. Biesmeijer, and R. Bommarco. 2012. Landscape context and elevation affect pollinator communities in intensive apple orchards. Basic and Applied Ecology 13(8):681–689. https://doi.org/10.1016/j.baae.2012.09.003

Mark D. M. 1984. Part 4: mathematical, algorithmic and data structure issues: automated detection of drainage networks from digital elevation models. Cartographica: The International Journal for Geographic Information and Geovisualization 21:168–178. https://doi.org/10.3138/10LM-4435-6310-251R

Matson P. A., W. J. Parton, A. G. Power, and M. J. Swift. 1997. Agricultural intensification and ecosystem properties. Science 277:504–509. https://doi.org/10.1126/science.277.5325.504

McRae B. H., B. G. Dickson, T. H. Keitt, and V. B. Shah. 2008. Using circuit theory to model connectivity in ecology, evolution, and conservation. Ecology 89(1):2712–2724. https://doi.org/10.1890/07-1861.1

McRae B. H., P. Beier. 2007. Circuit theory predicts gene flow in plant and animal populations. Proceedings of the National Academy of Sciences of the United States of America 104(50):19885–19890. https://doi.org/10.1073/pnas.0706568104

MEA (Millennium Ecosystem Assessment). 2005. Ecosystems and human well-being. Island Press, Washington, DC.

Melles S., N. Jones, B. Schmidt, and B. Rayfield. 2011. A least-cost path approach to stream delineation using lakes as patches and a digital elevation model as the cost surface. Procedia Environmental Sciences 7:240–245. https://doi.org/10.1016/j.proenv.2011.07.042

Melo D., B. Filgueiras, C. Iserhard, L. Iannuzzi, A. Freitas, and I. Leal. 2019. Effect of habitat loss and fragmentation on fruit-feeding butterflies in the Brazilian Atlantic Forest. Canadian journal of zoology 97(7):588–596. https://doi.org/10.1139/cjz-2018-0202

Middleton N., T. Sternberg. 2013. Climate hazards in drylands: A review. Earth-Science Reviews 126:48–57. https://doi.org/10.1016/j.earscirev.2013.07.008

Mills A., M. Fey. 2004. Effects of vegetation cover on the tendency of soil to crust in South Africa. Soil Use and Management 20(3):308–317. https://doi.org/10.1111/j.1475-2743.2004.tb00375.x

Mitchell M. G., E. M. Bennett, and A. Gonzalez. 2013. Linking landscape connectivity and ecosystem service provision: current knowledge and research gaps. Ecosystems 16:894–908. https://doi.org/10.1007/s10021-013-9647-2

Mitchell M. G., K. Chan, N. K. Newlands, and N. Ramankutty. 2020. Spatial Correlations Don’t Predict Changes in Agricultural Ecosystem Services: A Canada-Wide Case Study. Frontiers in Sustainable Food Systems 4:235. https://doi.org/10.3389/fsufs.2020.539892

Mitchell M. G., A. F. Suarez-Castro, M. Martinez-Harms, M. Maron, C. McAlpine, K. J. Gaston, K. Johansen, and J. R. Rhodes. 2015. Reframing landscape fragmentation’s effects on ecosystem services. Trends in Ecology & Evolution 30(4):190–198. https://doi.org/10.1016/j.tree.2015.01.011

Nassauer J. I., P. Opdam. 2008. Design in science: extending the landscape ecology paradigm. Landscape Ecology 23:633–644. https://doi.org/10.1007/s10980-008-9226-7

Neale T., J. Carmichael, and S. Cohen. 2007. Urban water futures: a multivariate analysis of population growth and climate change impacts on urban water demand in the Okanagan basin, BC. Canadian Water Resources Journal 32(4):315–330. https://doi.org/10.4296/cwrj3204315

Nelson E., G. Mendoza, J. Regetz, S. Polasky, H. Tallis, D. Cameron, K. M. Chan, G. C. Daily, J. Goldstein, and P. M. Kareiva. 2009. Modeling multiple ecosystem services, biodiversity conservation, commodity production, and tradeoffs at landscape scales. Frontiers in Ecology and the Environment 7(1):4–11. https://doi.org/10.1890/080023

Nicholls C. I., M. A. Altieri. 2013. Plant biodiversity enhances bees and other insect pollinators in agroecosystems. A review. Agronomy for Sustainable development 33(2):257–274. https://doi.org/10.1007/s13593-012-0092-y

Nicholson E., G. M. Mace, P. R. Armsworth, G. Atkinson, S. Buckle, T. Clements, R. M. Ewers, J. E. Fa, T. A. Gardner, and J. Gibbons. 2009. Priority research areas for ecosystem services in a changing world. Journal of Applied Ecology 46(6):1139–1144. https://doi.org/10.1111/j.1365-2664.2009.01716.x

Nilsson C., C. A. Reidy, M. Dynesius, and C. Revenga. 2005. Fragmentation and flow regulation of the world’s large river systems. Science 308:405–408. https://doi.org/10.1126/science.1107887

Pal S., S. Talukdar. 2018. Application of frequency ratio and logistic regression models for assessing physical wetland vulnerability in Punarbhaba river basin of Indo-Bangladesh. Human and Ecological Risk Assessment: An International Journal 24(5):1291–1311. https://doi.org/10.1080/10807039.2017.1411781

Pal S., P. Singha, K. Lepcha, S. Debanshi, S. Talukdar, and T. K. Saha. 2021. Proposing multicriteria decision based valuation of ecosystem services for fragmented landscape in mountainous environment. Remote Sensing Applications: Society and Environment 21:100454. https://doi.org/10.1016/j.rsase.2020.100454

Parks S. A., K. S. McKelvey, and M. K. Schwartz. 2013. Effects of weighting schemes on the identification of wildlife corridors generated with least-cost methods. Conservation Biology 27(1):145–154. https://doi.org/10.1111/j.1523-1739.2012.01929.x

Pebesma E., R. S. Bivand. 2005. Classes and methods for spatial data: the sp package. R news 5:9–13. https://cran.r-project.org/package=sp

Peng J., Y. Yang, Y. Liu, Y. Du, J. Meersmans, and S. Qiu. 2018. Linking ecosystem services and circuit theory to identify ecological security patterns. Science of the total environment 644:781–790. https://doi.org/10.1016/j.scitotenv.2018.06.292

Peron T., C. Comin, D. Amancio, L. da F Costa, F. Rodrigues, and J. Kurths. 2014. Correlations between climate network and relief data. Nonlinear Processes in Geophysics 21(6):1127–1132. https://doi.org/10.5194/npg-21-1127-2014

Phalan B., M. Onial, A. Balmford, and R. E. Green. 2011. Reconciling food production and biodiversity conservation: land sharing and land sparing compared. Science (New York, N.Y.) 333:1289–1291. https://doi.org/10.1126/science.1208742

Phillips J. D. 2013. Sources of spatial complexity in two coastal plain soil landscapes. Catena 111:98–103. https://doi.org/10.1016/j.catena.2013.07.003

Phillips J. D., W. Schwanghart, and T. Heckmann. 2015. Graph theory in the geosciences. Earth-Science Reviews 143:147–160. https://doi.org/10.1016/j.earscirev.2015.02.002

Phillips M. L., B. E. McNellis, M. F. Allen, and E. B. Allen. 2019. Differences in root phenology and water depletion by an invasive grass explains persistence in a Mediterranean ecosystem. American Journal of Botany 106(9):1210–1218. https://doi.org/10.1002/ajb2.1344

Phillips S. J., P. Williams, G. Midgley, and A. Archer. 2008. Optimizing dispersal corridors for the Cape Proteaceae using network flow. Ecological Applications 18:1200–1211. https://doi.org/10.1890/07-0507.1

Pinto N., T. H. Keitt. 2009. Beyond the least-cost path: evaluating corridor redundancy using a graph-theoretic approach. Landscape Ecology 24:253–266. https://doi.org/10.1007/s10980-008-9303-y

Poff N. L., J. D. Allan, M. B. Bain, J. R. Karr, K. L. Prestegaard, B. D. Richter, R. E. Sparks, and J. C. Stromberg. 1997. The natural flow regime. Bioscience 47(11):769–784. https://doi.org/10.2307/1313099

Potts S. G., J. C. Biesmeijer, C. Kremen, P. Neumann, O. Schweiger, and W. E. Kunin. 2010. Global pollinator declines: trends, impacts and drivers. Trends in ecology & evolution 25(6):345–353. https://doi.org/10.1016/j.tree.2010.01.007

Power A. G. 2010. Ecosystem services and agriculture: tradeoffs and synergies. Philosophical transactions of the royal society B: biological sciences 365(1554):2959–2971. https://doi.org/10.1098/rstb.2010.0143

Priess J. A., M. Mimler, A. Klein, S. Schwarze, T. Tscharntke, and I. Steffan-Dewenter. 2007. Linking deforestation scenarios to pollination services and economic returns in coffee agroforestry systems. Ecological Applications 17(2):407–417. https://doi.org/10.1890/05-1795

Qin K., J. Li, and X. Yang. 2015. Trade-off and synergy among ecosystem services in the Guanzhong-Tianshui Economic Region of China. International journal of environmental research and public health 12(11):14094–14113. https://doi.org/10.3390/ijerph121114094

Qiu J., M. G. Turner. 2013. Spatial interactions among ecosystem services in an urbanizing agricultural watershed. Proceedings of the National Academy of Sciences of the United States of America 110:12149–12154.

Queiroz C., M. Meacham, K. Richter, A. V. Norström, E. Andersson, J. Norberg, and G. Peterson. 2015. Mapping bundles of ecosystem services reveals distinct types of multifunctionality within a Swedish landscape. AMBIO 44:89–101. https://doi.org/10.1007/s13280-014-0601-0

R Development Core Team. 2013. R: A language and environment for statistical computing. R Foundation for Statistical Computing, Vienna, Austria. http://www.R-project.org/

Raudsepp-Hearne C., G. D. Peterson. 2016. Scale and ecosystem services: how do observation, management, and analysis shift with scale—lessons from Québec. Ecology and Society 21(3). https://www.jstor.org/stable/26269960

Rayfield B., M. Fortin, and A. Fall. 2011. Connectivity for conservation: a framework to classify network measures. Ecology 92(4):847–858. https://doi.org/10.1890/09-2190.1

Redhead J. W., G. D. Powney, B. A. Woodcock, and R. F. Pywell. 2020. Effects of future agricultural change scenarios on beneficial insects. Journal of environmental management 265:110550. https://doi.org/10.1016/j.jenvman.2020.110550

Renard D., J. M. Rhemtulla, and E. M. Bennett. 2015. Historical dynamics in ecosystem service bundles. Proceedings of the National Academy of Sciences of the United States of America 112(43):13411–13416. https://doi.org/10.1073/pnas.1502565112

Reynolds J. F., D. M. Smith, E. F. Lambin, B. L. Turner 2nd, M. Mortimore, S. P. Batterbury, T. E. Downing, H. Dowlatabadi, R. J. Fernandez, J. E. Herrick, E. Huber-Sannwald, H. Jiang, R. Leemans, T. Lynam, F. T. Maestre, M. Ayarza, and B. Walker. 2007. Global desertification: building a science for dryland development. Science 316:847–851. https://doi.org/10.1126/science.1131634

Ricketts T. H., J. Regetz, I. Steffan-Dewenter, S. A. Cunningham, C. Kremen, A. Bogdanski, B. Gemmill-Herren, S. S. Greenleaf, A. M. Klein, and M. M. Mayfield. 2008. Landscape effects on crop pollination services: are there general patterns? Ecology Letters 11(5):499–515. https://doi.org/10.1111/j.1461-0248.2008.01157.x

Rieb J. T., R. Chaplin-Kramer, G. C. Daily, P. R. Armsworth, K. Böhning-Gaese, A. Bonn, G. S. Cumming, F. Eigenbrod, V. Grimm, and B. M. Jackson. 2017. When, where, and how nature matters for ecosystem services: challenges for the next generation of ecosystem service models. Bioscience 67(9):820–833. https://doi.org/10.1093/biosci/bix075

Roa-García M., S. Brown, H. Schreier, and L. Lavkulich. 2011. The role of land use and soils in regulating water flow in small headwater catchments of the Andes. Water Resources Research 47(5). https://doi.org/10.1029/2010WR009582

Saha T. K., S. Pal. 2019. Emerging conflict between agriculture extension and physical existence of wetland in post-dam period in Atreyee River basin of Indo-Bangladesh. Environment, Development and Sustainability 21:1485–1505. https://doi.org/10.1007/s10668-018-0099-x

Satake A., T. K. Rudel, and A. Onuma. 2008. Scale mismatches and their ecological and economic effects on landscapes: A spatially explicit model. Global Environmental Change 18(4):768–775. https://doi.org/10.1016/j.gloenvcha.2008.07.007

Schlesinger W. H., J. F. Reynolds, G. L. Cunningham, L. F. Huenneke, W. M. Jarrell, R. A. Virginia, and W. G. Whitford. 1990. Biological feedbacks in global desertification. Science 247:1043–1048. https://doi.org/10.1126/science.247.4946.1043

Schröter M., A. Bonn, S. Klotz, R. Seppelt, and C. Baessler. 2019. Atlas of Ecosystem Services: Drivers, Risks, and Societal Responses. Springer, Cham.

Seavy N. E., T. Gardali, G. H. Golet, F. T. Griggs, C. A. Howell, R. Kelsey, S. L. Small, J. H. Viers, and J. F. Weigand. 2009. Why climate change makes riparian restoration more important than ever: recommendations for practice and research. Ecological Restoration 27(3):330–338. http://doi.org/10.3368/er.27.3.330

Simmonds J. S., B. J. van Rensburg, A. I. Tulloch, and M. Maron. 2019. Landscape-specific thresholds in the relationship between species richness and natural land cover. Journal of Applied Ecology 56(4):1019–1029. https://doi.org/10.1111/1365-2664.13320

Singleton P. H. 2002. Landscape permeability for large carnivores in Washington: a geographic information system weighted-distance and least-cost corridor assessment. US Department of Agriculture, Forest Service, Pacific Northwest Research Station, PNW-RP-549.

Stritih A., P. Bebi, and A. Grêt-Regamey. 2019. Quantifying uncertainties in earth observation-based ecosystem service assessments. Environmental modelling & software 111:300–310. https://doi.org/10.1016/j.envsoft.2018.09.005

Su C., B. Fu. 2013. Evolution of ecosystem services in the Chinese Loess Plateau under climatic and land use changes. Global and Planetary Change 101:119–128. https://doi.org/10.1016/j.gloplacha.2012.12.014

Suding K. N., S. Lavorel, F. Chapin Iii, J. H. Cornelissen, S. DIAz, E. Garnier, D. Goldberg, D. U. Hooper, S. T. Jackson, and M. Navas. 2008. Scaling environmental change through the community-level: A trait-based response- and-effect framework for plants. Global Change Biology 14(5):1125–1140. https://doi.org/10.1111/j.1365-2486.2008.01557.x

Tallis H., S. Polasky. 2009. Mapping and valuing ecosystem services as an approach for conservation and natural resource management. Annals of the New York Academy of Sciences 1162:265–283. https://doi.org/10.1111/j.1749-6632.2009.04152.x

Tallis H., P. Kareiva, M. Marvier, and A. Chang. 2008. An ecosystem services framework to support both practical conservation and economic development. Proceedings of the National Academy of Sciences of the United States of America 105(28):9457–9464. https://doi.org/10.1073/pnas.0705797105

Taylor P. D., L. Fahrig, K. Henein, and G. Merriam. 1993. Connectivity is a vital element of landscape structure. Oikos 68(3):571–573. https://doi.org/10.2307/3544927

Tilman D., K. G. Cassman, P. A. Matson, R. Naylor, and S. Polasky. 2002. Agricultural sustainability and intensive production practices. Nature 418:671–677. https://doi.org/10.1038/nature01014

Tomscha S. A., S. E. Gergel. 2016. Ecosystem service trade-offs and synergies misunderstood without landscape history. Ecology and Society 21(1):43. https://www.jstor.org/stable/26270345

Transboundary Connectivity Group. 2016. Providing a regional connectivity perspective to local connectivity conservation decisions in the British Columbia– Washington transboundary region: Okanagan-Kettle subregion connectivity assessment. Great Northern Landscape Conservation Cooperative.

Tscharntke T., R. Brandl. 2004. Plant-insect interactions in fragmented landscapes. Annual Reviews in Entomology 49:405–430. https://doi.org/10.1146/annurev.ento.49.061802.123339

Tscharntke T., A. M. Klein, A. Kruess, I. Steffan-Dewenter, and C. Thies. 2005. Landscape perspectives on agricultural intensification and biodiversity– ecosystem service management. Ecology Letters 8(8):857–874. https://doi.org/10.1111/j.1461-0248.2005.00782.x

Turner W. R., K. Brandon, T. M. Brooks, R. Costanza, G. A. Da Fonseca, and R. Portela. 2007. Global conservation of biodiversity and ecosystem services. Bioscience 57(1):868–873. https://doi.org/10.1641/B571009

Urban D. L., E. S. Minor, E. A. Treml, and R. S. Schick. 2009. Graph models of habitat mosaics. Ecology Letters 12(3):260–273. https://doi.org/10.1111/j.1461-0248.2008.01271.x

Van Berkel D. B., P. H. Verburg. 2014. Spatial quantification and valuation of cultural ecosystem services in an agricultural landscape. Ecological Indicators 37:163–174. https://doi.org/10.1016/j.ecolind.2012.06.025

Van der Ploeg S, De Groot RS, Wang Y. 2010. The TEEB Valuation Database: overview of structure, data and results. Wageningen, The Netherlands: Foundation for Sustainable Development. Foundation for Sustainable Development, Wageningen, Netherlands. NO DOI

van Etten J. 2017. R package gdistance: distances and routes on geographical grids. Journal of Statistical Software 76:1–21. https://doi.org/10.18637/jss.v076.i13

van Zanten B. T., I. Zasada, M. J. Koetse, F. Ungaro, K. Häfner, and P. H. Verburg. 2016. A comparative approach to assess the contribution of landscape features to aesthetic and recreational values in agricultural landscapes. Ecosystem Services 17:87–98. https://doi.org/10.1016/j.ecoser.2015.11.011

Verburg P. H., J. Van De Steeg, A. Veldkamp, and L. Willemen. 2009. From land cover change to land function dynamics: a major challenge to improve land characterization. Journal of environmental management 90(3):1327–1335. https://doi.org/10.1016/j.jenvman.2008.08.005

Wagner J. R. 2008. Landscape aesthetics, water, and settler colonialism in the Okanagan Valley of British Columbia. Journal of Ecological Anthropology 12(1):22–38. http://dx.doi.org/10.5038/2162-4593.12.1.2

Wagner J. R., K. White. 2009. Water and development in the Okanagan Valley of British Columbia. Journal of Enterprising Communities: People and Places in the Global Economy 3(4):378–392.

Wallace K. J. 2007. Classification of ecosystem services: problems and solutions. Biological Conservation 139(3-4):235–246. https://doi.org/10.1016/j.biocon.2007.07.015

Warman L. D., D. M. Forsyth, A. Sinclair, K. Freemark, H. D. Moore, T. W. Barrett, R. L. Pressey, and D. White. 2004. Species distributions, surrogacy, and important conservation regions in Canada. Ecology Letters 7:374–379.

Werling B. P., T. L. Dickson, R. Isaacs, H. Gaines, C. Gratton, K. L. Gross, H. Liere, C. M. Malmstrom, T. D. Meehan, L. Ruan, B. A. Robertson, G. P. Robertson, T. M. Schmidt, A. C. Schrotenboer, T. K. Teal, J. K. Wilson, and D. A. Landis. 2014. Perennial grasslands enhance biodiversity and multiple ecosystem services in bioenergy landscapes. Proceedings of the National Academy of Sciences of the United States of America 111(4):1652–1657. https://doi.org/10.1073/pnas.1309492111

Whittaker D., B. Shelby. 2002. Evaluating instream flows for recreation: Applying the structural norm approach to biophysical conditions. Leisure Sciences 24(3-4):363–374. https://doi.org/10.1080/01490400290050808

Wickham H. 2010. Stringr: modern, consistent string processing. The R Journal 2:38–40. https://doi.org/10.32614/RJ-2010-012

Xinxin L. 2017. Study on Spatial Heterogeneity of Urban Landscape Fragmentation in Changsha. Environmental Science and Management 1:9.

Zhang W., T. H. Ricketts, C. Kremen, K. Carney, and S. M. Swinton. 2007. Ecosystem services and dis-services to agriculture. Ecological Economics 64(2):253–260. https://doi.org/10.1016/j.ecolecon.2007.02.024

## Appendix References

Boerema A., A. J. Rebelo, M. B. Bodi, K. J. Esler, and P. Meire. 2017. Are ecosystem services adequately quantified? Journal of Applied Ecology 54(2):358–370. https://doi.org/10.1111/1365-2664.12696

Burkhard B., F. Kroll, S. Nedkov, and F. Müller. 2012. Mapping ecosystem service supply, demand and budgets. Ecological Indicators 21:17–29. https://doi.org/10.1016/j.ecolind.2011.06.019

ESRI. 2011. ArcGIS Desktop. Redlands, CA: Environmental Systems Research Institute. Release 10.

Gos P., S. Lavorel. 2012. Stakeholders’ expectations on ecosystem services affect the assessment of ecosystem services hotspots and their congruence with biodiversity. International Journal of Biodiversity Science, Ecosystem Services & Management 8(1-2):93–106. https://doi.org/10.1080/21513732.2011.646303

Hamel P., B. P. Bryant. 2017. Uncertainty assessment in ecosystem services analyses: seven challenges and practical responses. Ecosystem Services 24:1–15. https://doi.org/10.1016/j.ecoser.2016.12.008

Haughian S. R., P. J. Burton, S. W. Taylor, and C. Curry. 2012. Expected effects of climate change on forest disturbance regimes in British Columbia. BC Journal of Ecosystems and Management 13(1):1–24.

Hou Y., B. Li, F. Müller, Q. Fu, and W. Chen. 2018. A conservation decision-making framework based on ecosystem service hotspot and interaction analyses on multiple scales. Science of the Total Environment 643:277–291. https://doi.org/10.1016/j.scitotenv.2018.06.103

MoAg (BC Ministry of Agriculture). 2017. Agricultural Land Use Inventory (ALUI) dataset. Strengthening Farming, Abbostford, BC.

Okanagan Valley Economic Development Society (OVEDS). 2013. Economic Profile Okanagan Valley Kelowna, BC, Canada. Okanagan Valley Economic Development Society, Kelowna, BC.

Seppelt R., C. F. Dormann, F. V. Eppink, S. Lautenbach, and S. Schmidt. 2011. A quantitative review of ecosystem service studies: approaches, shortcomings and the road ahead. Journal of Applied Ecology 48(3):630–636. https://doi.org/10.1111/j.1365-2664.2010.01952.x

Wang Y., X. Li, Q. Zhang, J. Li, and X. Zhou. 2018. Projections of future land use changes: Multiple scenarios-based impacts analysis on ecosystem services for Wuhan city, China. Ecological Indicators 94(1):430–445. https://doi.org/10.1016/j.ecolind.2018.06.047

